# Population-specific effects of ocean acidification in the Olympia oyster

**DOI:** 10.1101/2023.09.08.556443

**Authors:** Laura H Spencer, Katherine Silliman, Steven Roberts

## Abstract

Populations of marine species that respond differently to ocean acidification offer natural reservoirs of biodiversity that can be leveraged for conservation efforts and to sustain marine food systems. The molecular and physiological traits associated with tolerance to acidification must first be identified. This study leveraged oysters from three phenotypically distinct populations of the Olympia oyster, *Ostrea lurida*, but that were bred and reared in common conditions for four years. We assessed their growth, reproductive development, and transcriptional response to acidification within and across generations. Responses reveal energetic trade-offs that reflect unique physiotypes previously observed among populations. The population with the slowest growth but high survival rates, oysters from Dabob Bay, mounted the largest transcriptional response to acidification without effects to growth and reproduction. A moderate response was observed in the population with fastest growth rate but lowest fecundity (Fidalgo Bay). Oyster Bay, the population with highest fecundity but lowest survival rates, did not respond at the transcript level. Oyster Bay was also the only population for which acidification negatively affected growth and reproductive development. While exposure to acidification did not affect gene expression in the next generation’s larval stage, it did result in larger larvae in the Oyster Bay population, which could partially alleviate negative effects of acidification in the wild for that population. Given the distinct transcriptional response of the Dabob Bay population to acidification and its high survival rates in previous studies, we then identified genes that were uniquely expressed in Dabob Bay oysters compared to the other populations. Genes involved in antibacterial and antiviral processes, metabolism, growth, and reproduction were uniquely expressed in Dabob Bay, and many similar functions were identified in both adults and larvae, which provides insight into the mechanisms behind a stress-tolerant oyster population. The population-specific physiotypes and responses to acidification illustrate the diversity of physiological strategies in *O. lurida* that balance the energetic demands of growth, reproduction, cellular maintenance, and offspring viability. Taken together this study reveals that there are distinct physiotypes among marine invertebrate populations on small geographic scales with implications for species resilience to acidification and other environmental stressors.

## Introduction

Following observations of shifting ocean conditions (IPCC, 2019) an enormous scientific effort has explored the response of marine species to ocean acidification (Riebesell & Gattuso, 2014). Empirical data has established that many species are vulnerable to ocean conditions projected for this century, particularly calcifying invertebrates, affecting a range of physiological processes over the lifetime of an organism, including development, recruitment, growth, reproduction, and survival (Gazeau et al., 2013; Kroeker et al., 2013; Lemasson et al., 2017; Melzner et al., 2019). However, these studies also indicate that biological responses are quite variable, related to an organism’s genetic and environmental ancestries (Eirin-Lopez & Putnam, 2019; He & Silliman, 2019; Przeslawski et al., 2015; Sunday et al., 2014). Some species are more tolerant to the effects of acidification than others (Branch et al., 2013; Figuerola et al., 2021), as are some populations within species (Bitter et al. 2019; Swezey et al. 2020; Kelly et al. 2013; Vargas et al. 2017). There is also evidence of intergenerational (spanning one generation) and transgenerational (spanning 2+ generations) plasticity, which may buffer future populations against challenging conditions (Salinas et al., 2013; Zhao et al., 2020). Ultimately, there will be a spectrum of responses to shifting ocean chemistry, dependent on species’ capacity to mitigate, acclimatize, and adapt to shifting conditions. For effective conservation and management of marine calcifiers, it is critical to identify the genotypes, physiotypes, and molecular mechanisms that impart tolerance to acidification, as well as quantify the range of responses within taxa.

Impacts of acidification to oysters were among the earliest observations of negative biological effects. Now, in an effort to build resilient commercial and wild stocks, breeding programs and researchers are increasingly seeking to identify oyster species and populations that are tolerant to acidification and other stressors. One such group of oysters appears to be species from the genus *Ostrea*, which includes the Olympia (*O. lurida*), European flat (*O. edulis*), Chilean flat (*O. chilensis*), and Australian flat oyster (*O. angasi*). Multiple studies have reported little to no effects of acidification in *Ostrea* spp. at the adult (Lemasson et al., 2018; Lemasson et al., 2019; Lemasson & Knights, 2021), juvenile (Navarro et al., 2020), and larval stages (Cole et al., 2016; Pereira et al., 2019; Waldbusser et al., 2016). In one study *O. edulis* larval growth and survival responded positively to acidification exposure (Prado et al., 2016). Unique *Ostrea* spp. life history traits, in particular brooding of veliger larvae in the maternal pallial cavity, may contribute to the species’ relative tolerance to acidification, as pH can quickly decrease to levels as low as 6.96 during periods of valve closures (Chaparro et al., 2009; Gray et al., 2019). There are, however, other studies that have observed negative effects of acidification in *Ostrea*, such as decreased larval growth in *O. lurida* that persist to the juvenile stage (Hettinger et al., 2012, 2013; Sanford et al., 2014). Contrasting effects could be explained by population-specific responses, which are commonly observed in marine invertebrate taxa (Barber et al., 1991; Macdonald & Thompson, 1988). Indeed, populations of *O. lurida* from northern California have diverged salinity tolerances, which is facilitated by distinct physiological responses at the cellular level (cell death regulation, mantle ciliary activity) (Maynard et al., 2018). Molecular strategies of *Ostrea* spp. need closer examination to understand the functions that enable tolerance to acidified conditions in some, but not all, populations (Melzner et al., 2009).

Previous studies have identified a suite of molecular functions in marine calcifiers that are sensitive to ocean acidification, which has been thoroughly reviewed by Melzner et al. (2019) and Strader et al. (2020). Interestingly, the directionality and magnitude of molecular responses can differ by species and/or populations within the same species. Opposing changes in antioxidants and molecular chaperones have been reported for acidification-tolerant (upregulated) and wild-type (downregulated) Sydney rock oysters in acidified conditions, which then reverse in subsequent generations (Goncalves et al., 2016, 2017). Changes to transcripts associated with energy production provide evidence for both metabolic depression and increased metabolic demand in response to acidification (Strader et al., 2020). In some cases the degree of transcriptional plasticity varies in response to acidification (Kenkel & Matz, 2017). For instance, upon exposure to acidification, transcripts for genes related to ATP production increased in two populations of urchins, but the magnitude of increase was more pronounced in the population that experiences more frequent periods of low pH compared to those from more stable pH environments (Evans et al., 2017). These studies highlight the variety of molecular responses to acidification that can occur among populations of the same species. It is vital to identify the fundamental molecular signatures of acidification-tolerant oysters to inform breeding and conservation efforts.

In the present study we examine effects of acidification on multiple populations of the Olympia oyster, *Ostrea lurida*. *O. lurida* is native to the North American Pacific Coast, inhabiting dynamic estuarine environments that are influenced by coastal upwelling and ocean acidification (Feely et al., 2010; McGraw, 2009; Reum et al., 2014). We build upon previous studies that have identified unique fitness traits in three populations from disparate regions of greater Puget Sound in Washington State (Figure 1, Table 1) (Heare et al., 2017; Heare et al., 2018; Silliman et al., 2018; Spencer et al., 2020; White et al., 2017). Oysters derived from Oyster Bay in the South Puget Sound are highly fecund, and compared to other populations require fewer degree-days to begin reproducing (Heare et al., 2017; Silliman et al., 2018; Spencer et al., 2020). Oysters from Dabob Bay in the Hood Canal basin consistently display slower growth than other populations but have higher survival rates in field testing and during hatchery rearing (Heare et al., 2017; Spencer et al., 2020, Ryan Crim*, pers. comm.*). Those from Fidalgo Bay, a small basin in the northern reaches of greater Puget Sound, grow faster and are less fecund or have delayed reproduction (Heare et al., 2017; Silliman et al., 2018; Spencer et al., 2020).

**Figure 1:**
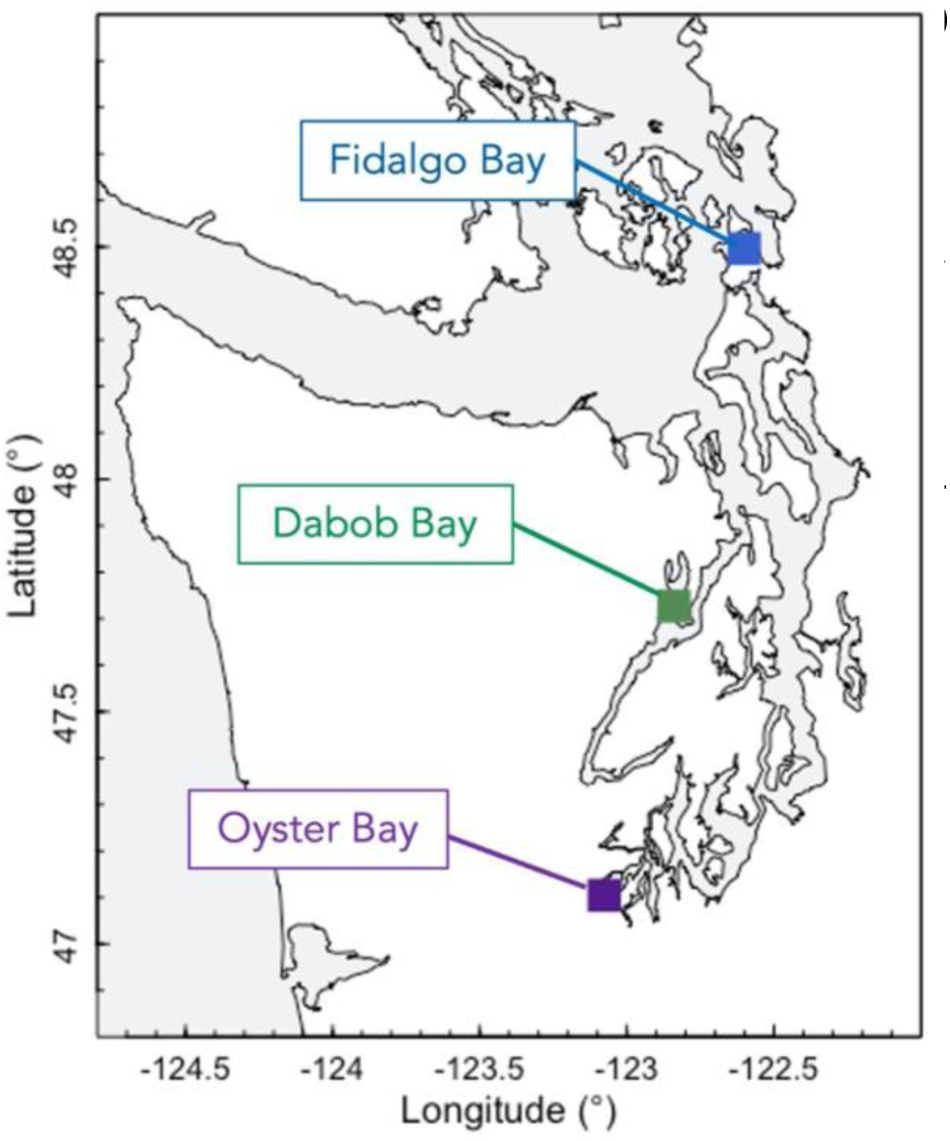
Location of the focal *O. lurida* populations in the greater Puget Sound estuary, Washington, USA.

**Table 1.** Traits that are characteristic of the focal *O. lurida* populations from the greater Puget Sound region, summarized from previous studies of the same populations: (1) Heare et al. 2017 (2) Heare et al. 2018 (3) Silliman et al. 2018 (4) Spencer et al. 2020 (5) Ryan Crim *pers. comm*. Additionally, genetic differentiation among populations was characterized by White et al. (2017).

| Location of *O. lurida* population | Growth rate, size at maturity | Reproductive output, timing | Performance in field & hatchery trials | Immediate transcriptional response to stress |
| --- | --- | --- | --- | --- |
| Fidalgo Bay | Fast growth, large at maturity<sup>1,3</sup> | Low/Moderate fecundity<sup>1,3,4</sup> | Moderate survival<sup>1,4</sup> | No response<sup>2</sup> |
| Dabob Bay | Slow growth, small at maturity<sup>1,3,4</sup> | Moderate fecundity<sup>1,3,4</sup> | High survival in field, high larval survival in hatchery<sup>1,4,5</sup> | No response<sup>2</sup> |
| Oyster Bay | Moderate growth, medium/large at maturity<sup>1,3,4</sup> | Very high fecundity, early to reproduce<sup>1,3,4</sup> | Low/moderate survival in field<sup>1,4</sup> | Altered expression of targeted genes 1-hr following heat & mechanical stress<sup>2</sup> |

Here, we use high-throughput sequencing to perform the first transcriptional characterization of an *Ostrea* species to ocean acidification. Expression analyses are paired with biometric data (gonad development, growth) to capture system-wide changes in energy allocation due to acidification exposure (Sokolova et al., 2012). By examining oysters from the aforementioned Olympia oyster populations (Dabob Bay, Fidalgo Bay, and Oyster Bay) we leverage a comprehensive understanding of the traits characteristic of each population (Table 1) (Heare et al., 2017; Heare et al., 2018; Silliman et al., 2018; Spencer et al., 2020), and by using individuals that were bred and grown in common conditions we control for within-generation carryover effects (Hettinger et al., 2012, 2013). Given the possible influence of intergenerational exposures on an organism’s physiology (Goncalves et al., 2016, 2017), we also extend the analysis to include a second generation (larval offspring) to examine the potential impacts of parental exposures and population-of-origin on basal functions (larval size, gene expression). By fully describing its molecular and physiological response to acidification across distinct populations we show that *O. lurida* is a good candidate for aquaculture investment & conservation.

## Methods

### Adult oyster source and history

Experimental oysters were bred in 2013 as described in Heare et al. (2017) in common conditions from three populations of wild broodstock, which were harvested from Dabob Bay in Hood Canal, Oyster Bay in South Puget Sound, and Fidalgo Bay in North Puget Sound. Oysters were then maintained in common conditions in a pearl net adjacent to the Kenneth K. Chew Center for Shellfish Research and Restoration in central Puget Sound for approximately 3.5 years. Upon entering experimental conditions, shell height for each population was on average 29.8 ± 4.6mm, 35.7 ± 4.5mm, and 35.7 ± 4.4mm for Dabob Bay, Oyster Bay, and Fidalgo Bay, respectively. Experimental oysters were therefore mature adults that had been produced and reared in common conditions, but had distinct genetic heritage.

### Adult treatments and larval collection

Beginning February 16, 2017 adults were exposed to two pCO_2_ treatments for 52 days (control pCO_2_: 841 ± 85 μatm, pH 7.82 ± 0.02; high pCO_2_: 3,045 ± 488 μatm, pH 7.31 ± 0.02), as described in Spencer et al. (2020) and Venkataraman et al. (2019) (Supplemental Materials). Following experimental exposures, adults from all populations and pCO_2_ treatments were returned to common conditions, reproductively conditioned, and spawned to produce offspring. Because *O. lurida* are viviparous spermcasters and brood larvae to the veliger stage, larvae were captured upon maternal liberation, which commenced on May 14th and persisted for 57 days. Details of the experimental and spawning conditions are described in (Spencer et al., 2020). Prior to the pCO_2_ treatments adults were also exposed to two temperature regimes for 60 days. These represent what would be considered normal or ambient temperature (6.1 ± 0.2°C) and a temperature that would be considered elevated at the experimental site (10.2 ± 0.5°C) (see Spencer et al. 2020). Given that temperature did not interact with pCO_2_ to affect the focal characteristics in adults or larvae, only effects of parental pCO_2_ exposure were further examined for this study.

### Tissue sampling

Adult ctenidia tissue was collected immediately upon terminating pCO_2_ treatments from Dabob Bay, Fidalgo Bay, and Oyster Bay. Nine oysters from each population and pCO_2_ treatment were sacrificed and ctenidia tissue was collected and flash-frozen at approximately −116°C using a solution of ethanol and dry-ice then preserved at −80°C. Gonad tissue was collected from the same individuals, preserved in histology cassettes using the PAXgene Tissue FIX System (PreAnalytiX, Hombrechtikon, Switzerland), then processed for reproductive development analysis by Diagnostic Pathology Medical Group (Sacramento, California, USA).

Larval offspring were sampled at the veliger stage upon maternal liberation. For each group of larvae that was released, a portion were reared as described in Spencer et al. (2020) and the remaining were preserved by rinsing with fresh water into microcentrifuge tubes, removing water, then placing directly into −80°C freezer. Given the variable reproductive rates, the number of larval groups preserved for downstream analysis from control pCO_2_ and high pCO_2_ varied, and was 7 and 10 for Fidalgo Bay, 5 and 7 for Dabob Bay, and 17 and 13 for Oyster Bay, respectively. Each larval sample contained thousands of larvae (on average 170k), and consisted of siblings that were released from the same female on the same day. However, given that adults were continuously spawning and releasing larvae, it is possible that some larval samples contained a mix of multiple families from the same population and treatment.

### Adult growth and reproductive development

Adult oysters were measured for shell height before and after pCO_2_ treatments (n=9 per population x pCO_2_ treatment) using digital calipers (mm), defined as the maximum distance from the umbo along the dorsal/ventral axis. Shell height after pCO_2_ treatments were compared among population and pCO_2_ treatments using 2-way ANOVA. Since we were most interested in assessing effects of high pCO_2_ exposure on growth rate for each population, population-specific differences in size between high-, control- and pre-pCO_2_ treatments were tested using 1-way ANOVA, and Tukey honest significant difference tests were performed to assess pairwise comparisons.

Gonad samples (n=9 per population x pCO_2_ treatment) collected before and after pCO_2_ treatments were assigned sex and stage as described in Spencer et al, (2020). For each population, contingency tables were constructed for gonad sex, developmental stage of sperm, and developmental stage of eggs, and differences between pCO_2_ treatments were compared using Chi-Square or Fisher Exact Tests (depending on the number of males or females present within a population) and P-values were estimated using Monte-Carlo simulations with 1,000 permutations.

### Larval offspring size

Newly liberated veliger larvae were measured using a Nikon eclipse Ni microscope and the NIS-Elements BR imaging and measuring software (version 4.60). Mean shell height (distance from hinge to margin, perpendicular to hinge), and mean shell width (longest distance parallel to hinge) were estimated from at minimum 48 larvae per collection from each tank. Mean shell height and width were compared among population and parental treatments using 2-Way ANOVA and Type II Sums of Squares with the car package (Fox & Weisberg, 2018). Then, since we were most interested in assessing population-specific responses, effects of parental pCO_2_ on larval size were assessed for each population using 1-Way ANOVA with Type II Sums of Squares.

### Library construction and sequencing

RNA was isolated from 52 frozen adult ctenidia samples (n=8 or 9 per population x pCO_2_ treatment) and 61 pooled whole-body larvae (5-13 per population x parental pCO_2_ treatment) (Table 2). Each sample was homogenized in liquid nitrogen with a stone mortar and pestle and isolated following the RNAzol® RT protocol for Total RNA Isolation (Molecular Research Center, Inc., Cincinnati, OH). RNA pellets were resuspended in DEPC-treated water, residual DNA contamination was removed using the Turbo DNase kit (Life Technologies, Carlsbad, CA), then RNA was quantified using the Qubit RNA assay with Qubit 3.0 Fluorometer (2.0ul of each sample for quantification, Life Technologies, Carlsbad, CA), and quality was assessed for a subset of RNA isolates using the Bioanalyzer RNA 6000 Pico Chip assay (Agilent Technologies, Santa Clara, CA).

**Table 2.**
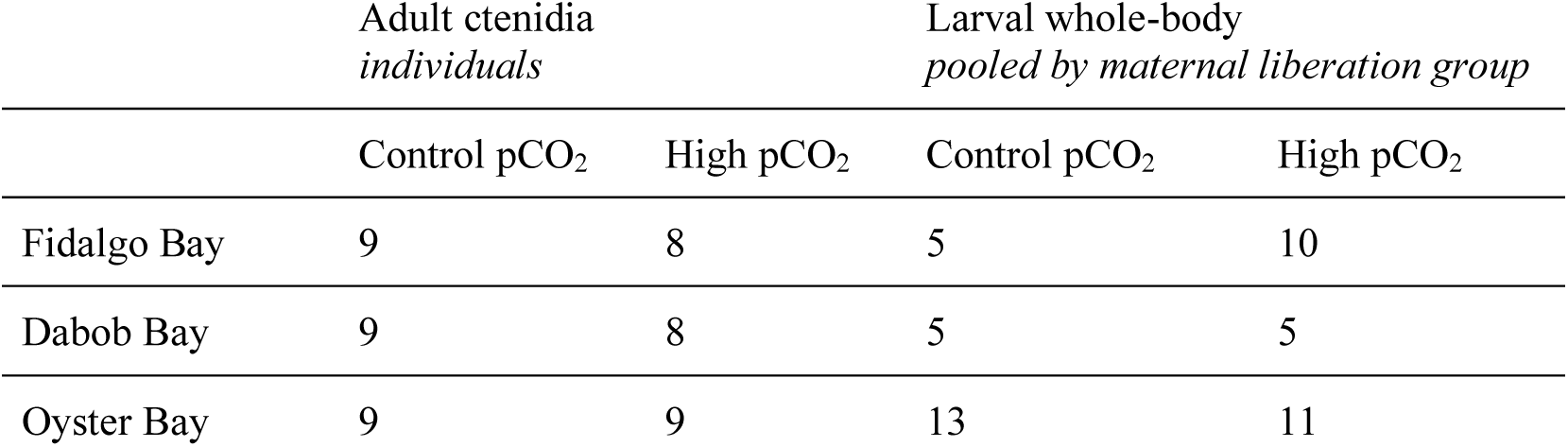
The number of QuantSeq libraries examined per population and parental pCO_2_ treatment for adult and larval tissues.

|  | Adult ctenidia<br>*individuals* |  | Larval whole-body<br>*pooled by maternal liberation group* |  |
| --- | --- | --- | --- | --- |
|  | Control pCO<sub>2</sub> | High pCO<sub>2</sub> | Control pCO<sub>2</sub> | High pCO<sub>2</sub> |
| Fidalgo Bay | 9 | 8 | 5 | 10 |
| Dabob Bay | 9 | 8 | 5 | 5 |
| Oyster Bay | 9 | 9 | 13 | 11 |

Library preparation was performed following the QuantSeq 3’ mRNA-Seq protocol (v.015UG009V0251, Lexogen, Vienna, Austria), also known as TagSeq, which generates cDNA from the 3’ end of mRNA strands and only generates one fragment per mRNA transcript, allowing for accurate gene expression data at a reduced cost (Meyer et al., 2011). Briefly, Total RNA (350 ng) was used to generate single-stranded DNA using reverse transcription (oligoT priming), which binds to the poly(A) tail and includes the read 1 adapter; RNA was removed, then the second DNA strand + Illumina adapter was synthesized by random priming; the double stranded cDNA + adapters were purified using magnetic beads, then an aliquot (1uL) of each library was amplified using qPCR to determine the optimal number of cycles needed for Endpoint PCR, which ranged from 14 to 17; the libraries were then amplified and indexed with unique barcodes, then re-purified. Prior to sequencing, all libraries were quantified using Qubit High Sensitivity DNA kits (Life Technologies, Carlsbad, CA), and the quality (e.g. fragment length) was assessed for a subset using a Bioanalyzer High Sensitivity DNA chip kit (Agilent Technologies, Santa Clara, CA). Single-end sequencing with 100-bp read length was conducted on a NovaSeq platform (Illumina, San Diego, CA) by the University of Washington’s Northwest Genomics Center, who also demultiplexed the raw sequencing data.

### Expression data processing & analysis

Raw reads were inspected for quality using FastQC (Andrews 2010) and MultiQC (Ewels et al., 2016), trimmed to remove Illumnina adapters, poly(A)- and poly(G)-tails, and quality filtered (≥20 read length, >15 quality score) using the Cutadapt toolkit v2.10 (Martin, 2011), then aligned to the draft *O. lurida* genome v081 (GenBank accession GCA_903981925.1) using Bowtie2 v2.4.1 (Langmead & Salzberg, 2012) and local alignment with the pre-set --sensitive-local option. Average alignment rate was 75.7±4.4% for adult libraries and 70.7±8.2% for larval libraries. From the resulting alignment files (.bam), the number of reads that uniquely mapped to each *O. lurida* gene was determined using featureCounts v2.0.0 (Liao et al., 2014) and a gene annotation file that was adjusted to extend the 3’ end by 2kb, as libraries were generated at the 3’ end of the original mRNA transcripts.

Differential gene expression analysis was performed using DESeq2 (Love et al., 2014) in R v4.0.4 using RStudio interface v1.3.1093 (R Core Team, 2021; RStudio Team, 2020). Read counts were first filtered to remove genes with fewer than 10 total counts across all samples (1,229 and 7,725 genes were discarded for adult ctenidia and larval pools, respectively). For adults the number of reads retained for analysis ranged from 1.18M to 3.30M per sample and averaged 1.99M ±SD442K, and for larvae total reads ranged from 653K to 2.04M and averaged 1.32M ±SD332K. The total number of genes and per-sample average retained for analysis was 30,981 and 28,110±607 for adult ctenidia, and 24,485 and 23,114 ±SD1,259 for larvae. Gene counts were assessed for differential gene expression for adult ctenidia between pCO_2_ exposure within populations, and for larvae between parental pCO_2_ exposures within and across populations. DESeq2 uses raw count data to generate generalized linear models and internally corrects for library size, therefore counts were not normalized prior to differential expression analysis.

Given the high performance of the Dabob Bay population in previous lab and field trials (Heare et al. 2017; Spencer et al. 2020; Ryan Crim *pers. comm.*), we examined genes that were constitutively expressed at unique levels in Dabob Bay. Pairwise differences in expression among Dabob Bay and the other two populations were identified in DESeq2 using transcriptomes from adults held in control conditions (n=9 libraries per population), and separately using transcriptomes from all larval samples (Table 2). For each life stage (adult and larvae), genes from the two pairwise comparisons (Dabob vs Fidalgo, Dabob vs. Oyster) were clustered into two gene sets: 1) genes that were more abundant, and 2) genes that were less abundant than the other two populations. Overlapping genes uniquely abundant in both adults and larvae from Dabob Bay were then interrogated for relative expression patterns (i.e. higher or lower than the other populations).

Differentially expressed genes were merged with the *O. lurida* genome to generate lists of Uniprot IDs from annotated genes. Enriched biological processes of differentially expressed genes sets were determined using the Gene-Enrichment and Functional Annotation Tool from DAVID v6.8, and were defined as those with modified Fisher Exact P-Values (EASE Scores) <0.1.

## Results

### Effects of acidification on adult growth, sex ratio

Adult oyster growth rate was reduced by acidification in the Oyster Bay population only: shell size did not differ after the high pCO_2_ exposure, whereas those that were exposed to control pCO_2_ were larger after treatment (Figure 2, Table 3). The Dabob Bay and Fidalgo Bay populations did not differ in size after either pCO_2_ treatment compared to before (Figure 2, Table 3). Shell height differed by population before (F(2,72)=3.44, p=0.034) and after (F(2,84)=16.5, p=9.28e^-7^) pCO_2_ treatments, with the Dabob Bay population significantly smaller than both Fidalgo Bay and Oyster Bay, particularly after pCO_2_ treatments.

**Figure 2.**
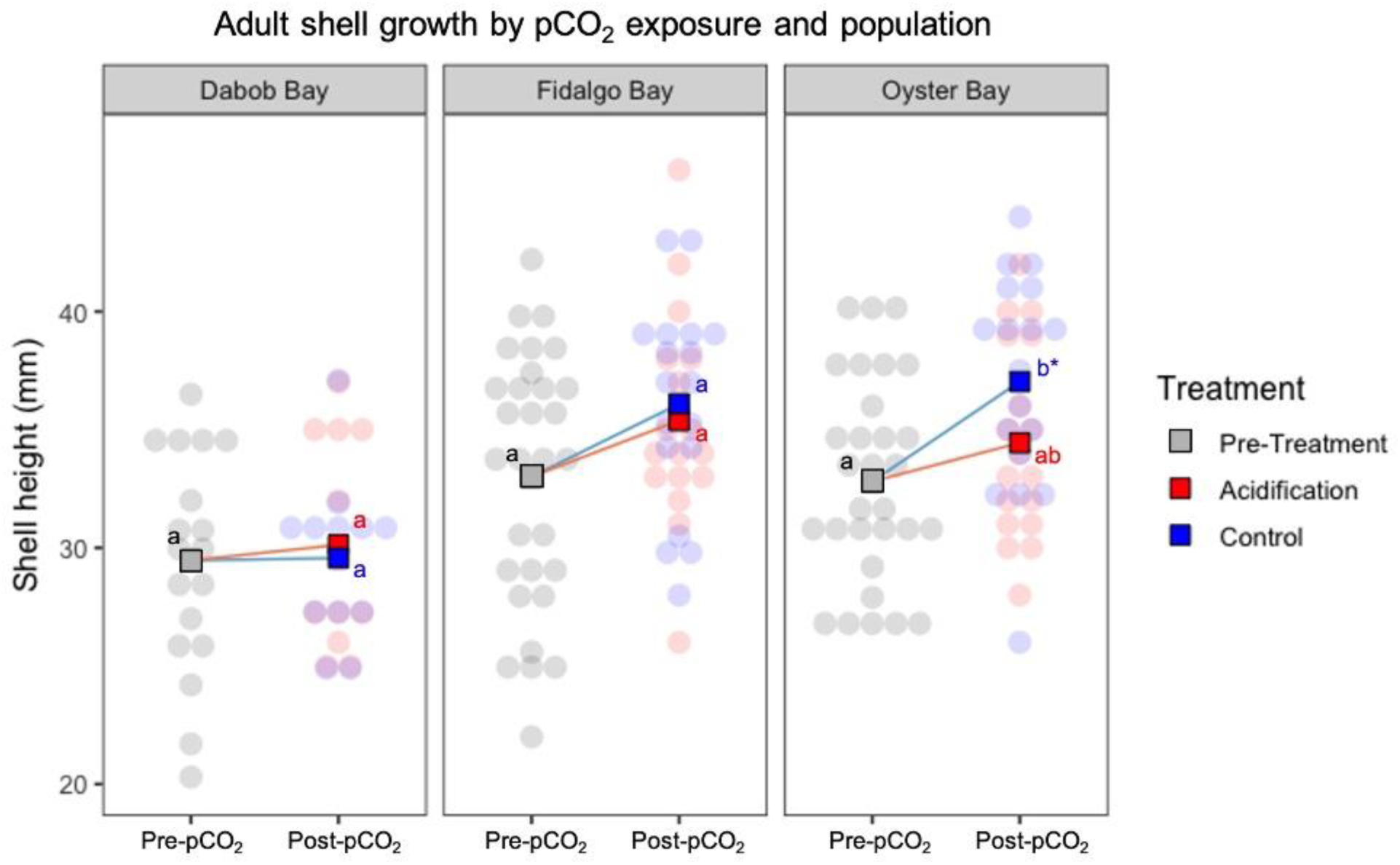
Adult shell height before pCO_2_ treatment (“Pre-pCO_2_”) and after 52 days in acidification (pCO_2_ = 3045 ± 488 μatm: pH = 7.31 ± 0.02) and control conditions (pCO_2_ = 841 ± 85 μatm; pH = 7.82 ± 0.02) for each population. Transparent points each represent one adult oyster color-coded by pCO_2_ treatment, and overlaid square points indicate mean shell height. Those exposed to control pCO_2_ were larger than before the pCO_2_ treatments in the Oyster Bay population only.

**Table 3:** Shell size statistics for each population. 1-way ANOVA compared shell height of oysters before treatment (Pre-Treatment) and after acidification and control treatments. Tukey multiple comparison of means was used to test for significant growth of oysters during the 52-day exposure to control conditions and acidification.

| Population | ANOVA statistics | Tukey comparison of means:<br>*Control vs. Pre-treatment* | Tukey comparison of means:<br>*Acidification vs. Pre-treatment* |
| --- | --- | --- | --- |
| Dabob Bay | F(2,39)=0.09, p=0.91 | $\Delta=0.10$, p-adj=1.0 | $\Delta=0.66$, p-adj=0.91 |
| Fidalgo Bay | F(2,63)=0.26, p=0.079 | $\Delta=3.08$, p-adj=0.094 | $\Delta=2.38$, p-adj=0.24 |
| Oyster Bay | **F(2,63)=5.35, p=0.0071\*** | **$\Delta=4.22$, p-adj=0.0049\*** | $\Delta=1.64$, p-adj=0.42 |

Progression from one sex to the other in these hermaphroditic oysters was affected by acidification in the Oyster Bay population only: the ratio of predominantly-females:predominantly-males differed significantly between pCO_2_ treatments (p-sim=0.015), with fewer females present after exposure to high pCO_2_ (Figure 3). The developmental stages of sperm and eggs (when present) did not differ between control and high pCO2 treatment in any population. Sperm development did not change during the 52-day exposure period in any population. The Fidalgo Bay population exposed to control conditions contained more advanced- and late-stage oocytes following the 52-day exposure compared to before (p-sim=1.0e-4), but this was not the case for Fidalgo Bay oysters exposed to acidification, or for any other population. Additionally, compared to before pCO_2_ treatment, the sex ratio differed following control pCO_2_ exposure in Oyster Bay (p-sim= 8.0e^-4^) and Fidalgo Bay (p-sim=1.3e^-3^), but did not differ following high pCO_2_ exposure in any population, or between any comparison in the Dabob Bay population.

**Figure 3.**
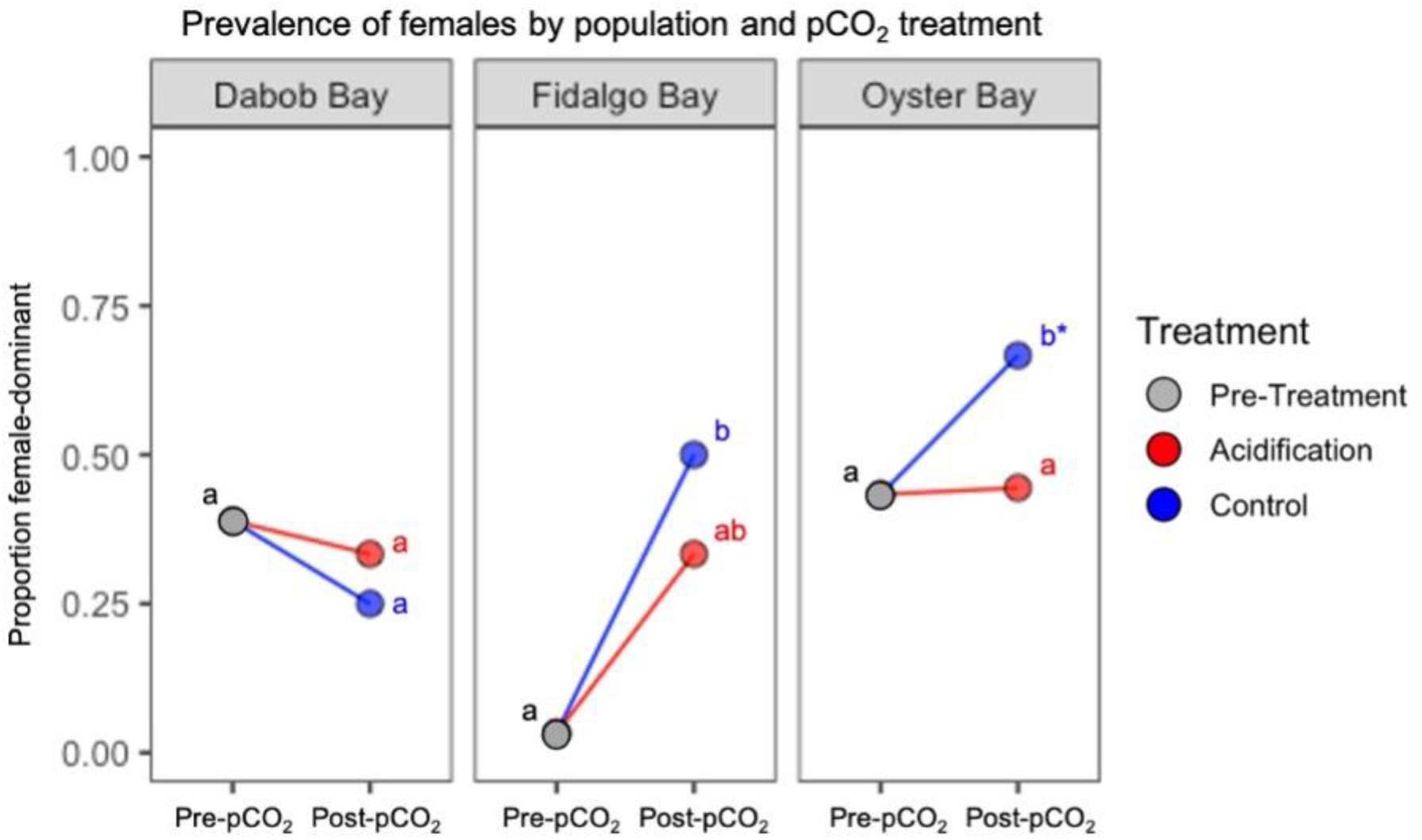
Proportion of oysters that were female or female-dominant before pCO_2_ treatment (“Pre-pCO_2_”) and after 52 days in acidification (pCO_2_ = 3045 ± 488 μatm: pH = 7.31 ± 0.02) and control conditions (pCO_2_ = 841 ± 85 μatm; pH = 7.82 ± 0.02) for each population. The prevalence of females differed among pCO_2_ treatments in the Oyster Bay population only.

The developmental stages of sperm and eggs (when present) did not differ between control and high pCO_2_ treatment in any population. Sperm development did not change during the 52-day exposure period in any population. The Fidalgo Bay population exposed to control conditions contained more advanced- and late-stage oocytes following the 52-day exposure compared to before (p-sim=1.0e^-4^), but this was not the case for Fidalgo Bay oysters exposed to acidification, or for any other population.

### Differential expression in adults upon direct exposure to acidification

Of 32,210 genes in the draft *O. lurida* genome, we detected expression in 30,981 genes in adult ctenidia tissue. Within populations, 132 and 76 genes were differentially expressed in Dabob Bay and Fidalgo Bay in response to pCO_2_ treatments, respectively (Supplemental Materials). No expression differences were detected in the Oyster Bay population upon exposure to pCO_2_ treatments. The annotated Dabob Bay and Fidalgo Bay DEGs were enriched for 25 and 6 biological functions (Figure 4). Four genes were differentially expressed in both Dabob Bay and Fidalgo Bay, which code for Cytochrome P450 2B4 (CYP2B4, p-adj=0.021), Cytochrome P450 2U1 (CYP2U1, p-adj=0.015), Fatty acid-binding protein (FABP4, p-adj=0.032), and Thyroxine 5-deiodinase (DIO3, p-adj=0.028), all of which were more abundant in oysters exposed to high pCO_2_.

**Figure 4.**
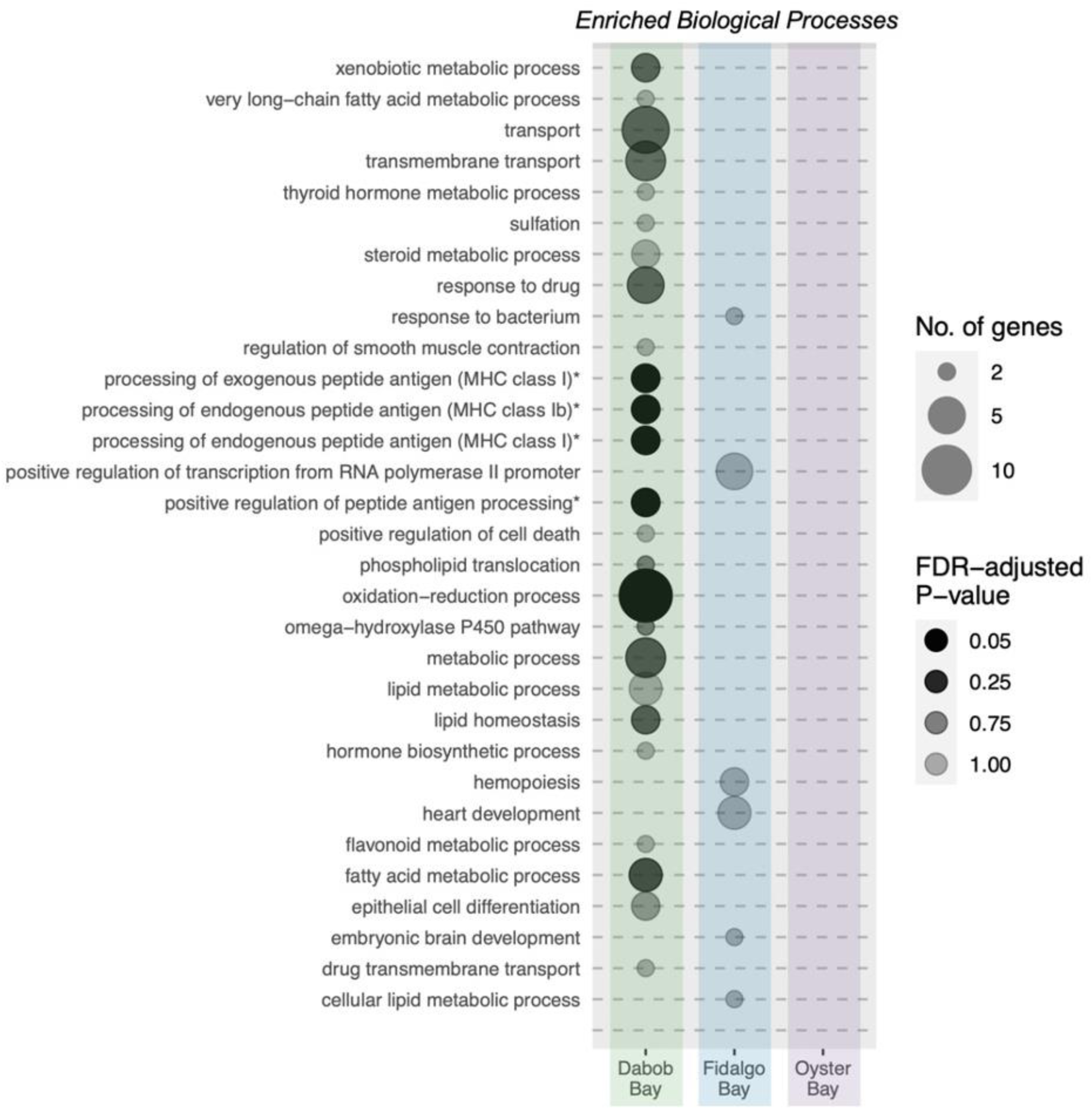
Enriched biological processes of differentially expressed genes upon exposure to high pCO_2_ in three populations with different bays-of-origin. No genes were differentially expressed in the Oyster Bay population, and thus no processes were enriched. Asterisks indicate GO terms that have been edited for length.

### Impacts of parental exposures on larval offspring size

Within populations, larval shell width and height differed by parental pCO_2_ in the Oyster Bay population only (width: F(1,29)=5.46, p=0.027; height: F(1,29)=4.56, p=0.041), but did not differ significantly among parental pCO_2_ in Fidalgo Bay (width: F(1,15)=0.0071, p=0.93; height: F(1,159)=0.14, p=0.71), or Dabob Bay (width: F(1,10)=0.71, p=0.49; height: F(1,10)=0.10, p=0.76) (Figure 5).

**Figure 5.**
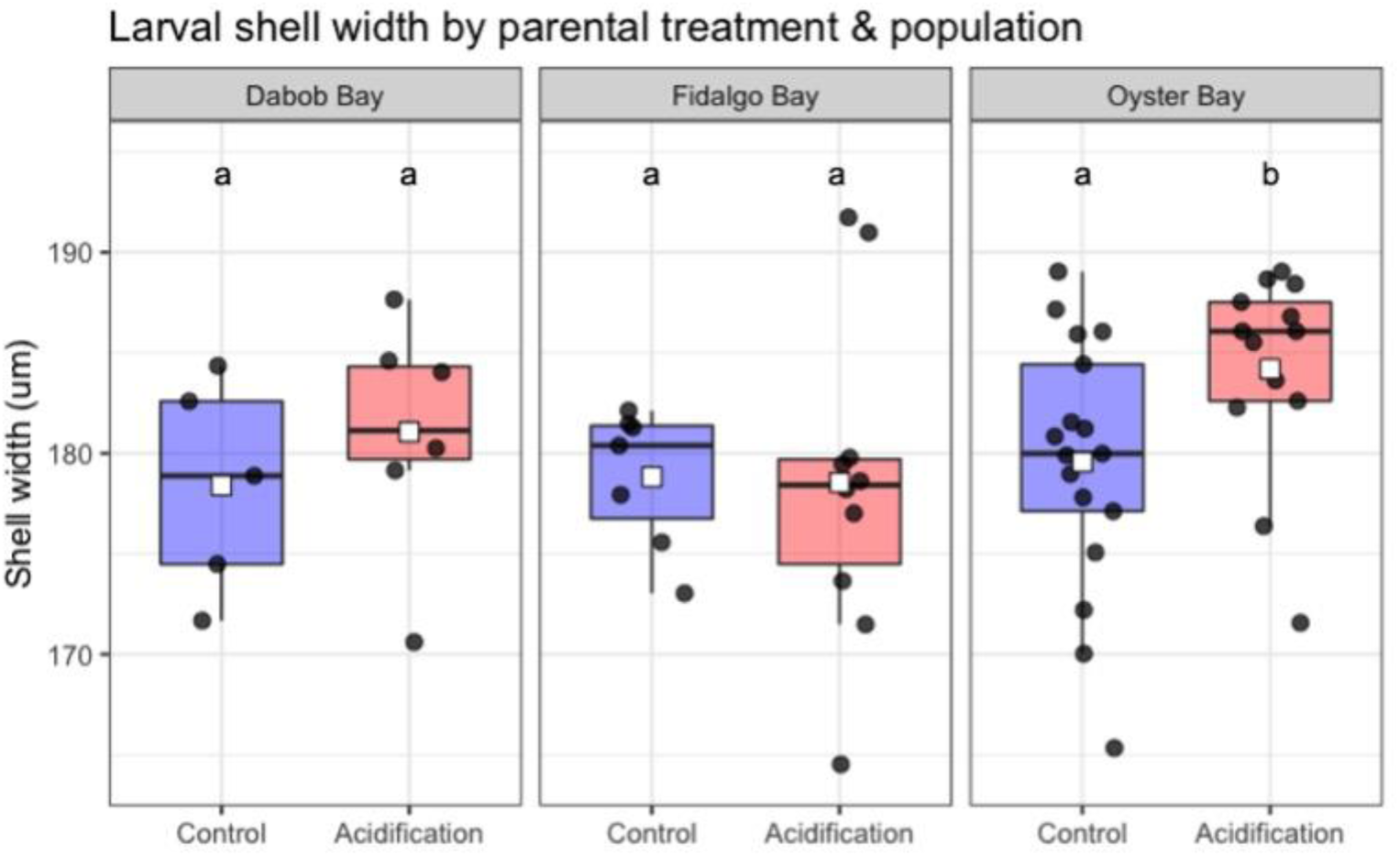
Laval shell size upon maternal liberation was significantly affected by parental exposure to high pCO_2_ in the Oyster Bay population only. Shell width is shown here. Each point represents the mean shell width of 40+ larvae from a release group. Treatments were terminated prior to adult reproductive conditioning and spawning, so larvae were never directly exposed to treatments.

### Larval offspring gene expression

Larvae were pooled by maternal release pulse and assessed for gene expression differences by parental exposure to acidification. We detected expression of 24,485 genes in larval samples. Within populations, one gene was differentially expressed by parental pCO_2_ in the Fidalgo Bay population, “OLUR_00020618” (log2 fold change = −1.6, p-adj = 0.011), which was not annotated. Across all populations no expression differences were detected between parental pCO_2_ exposures.

### Constitutive expression differences in the high-performing Dabob Bay population

Of the 30,981 genes examined in adults, 280 differed between Dabob Bay and Oyster Bay (0.9% of genes), and 379 differed between Dabob Bay and Fidalgo Bay (1.2% of genes), 89 of which were identified by both comparisons. Of these genes uniquely expressed in Dabob Bay, 220 were annotated (Supplemental Materials) and were enriched for 16 biological processes. There were 31 genes that were both constitutively expressed at different levels in Dabob Bay compared to other populations and were differentially expressed in response to acidification. Of these overlapping genes, eleven were less abundant than the other populations constitutively, but in response to acidification they increased in abundance. Twenty were more abundant constitutively, but in response to acidification they became less abundant.

Of the 24,485 genes examined in larval offspring, 260 and 452 differed between Dabob Bay and Fidalgo Bay and Oyster Bay, respectively (0.86% and 0.88%), 59 of which were identified by both comparison and 25 of which were annotated (Supplemental Materials). Forty-two genes were differentially expressed in both adult and larval stages among Dabob Bay and the other populations, which largely followed the same constitutive expression patterns in both life stages: 26 genes were more abundant in Dabob Bay adults and larvae compared to the other populations, and 14 were less abundant in both stages.

## Discussion

This study explored the within- and intergenerational response of *O. lurida* to acidification in three populations that had been reared in common conditions, but had distinct genetic ancestries. Each population demonstrated unique physiologies, which were evident in growth, gonad development, and transcriptional responses to pCO_2_ treatments. In the Dabob Bay population there was no growth or change in sex ratio regardless of treatment. However, Dabob Bay oysters demonstrated a robust transcriptional response to high pCO_2_ enriched for oxidation-reduction/detoxification, lipid metabolism, and other key processes. In stark contrast, no transcriptional response was detected in Oyster Bay oysters, but growth rate and prevalence of females were both negatively affected by acidification. There was a moderate transcriptional response in the Fidalgo Bay population, with no change in growth rate and minor effects to sex ratio as a result of acidification. This study also explored intergenerational carryover effects of adult exposure to acidification on basal gene expression and size of larvae upon maternal liberation across populations. Larvae were larger from adults exposed to high pCO_2_ in the Oyster Bay population only. Counter to predictions we found no signature of parental exposure to acidification in larval transcriptomes. The unique responses to acidification support previous observations of varying stress-tolerance in the same populations.

### Population-specific effects of acidification on growth and reproduction are associated with performance

Growth rate varies among Puget Sound *O. lurida* populations (Heare et al., 2017; Heare et al., 2018; Silliman et al., 2018; Spencer et al., 2020), and this study indicates that it also responds to acidification in a population-specific manner. Shell growth was stunted by acidification in the adults from Oyster Bay, as they grew in control conditions but did not in acidified conditions. In contrast, adults from Dabob Bay and Fidalgo Bay did not grow during the 52-day experiment regardless of pCO_2_ treatment. Previous studies characterized Dabob Bay as the slowest growing population, but also found that those oysters also performed best in hatchery and field trials. Heare et al. (2017) and Silliman et al. (2018) both observed slowest growth in the Dabob Bay population and fastest growth in the Fidalgo Bay population. Heare et al. (2017) also observed highest deployment survival in Dabob Bay progeny. In Spencer et al. 2020 the Dabob Bay population was slowest to reach the eyed larval stage (∼18 days) (Fidalgo Bay was the fastest, ∼14 days), and had the highest survival during the larval stage (the lowest survival was in Oyster Bay population). Together, these studies indicate that *O. lurid*a from Dabob Bay may prioritize stress resilience and survival at the cost of growth rate and size at maturity.

Acidification affected reproductive processes in a population-specific manner. The natural male-female sexual progression was significantly altered by acidification in the Oyster Bay population only. Reproductive traits appear to be heritable in Puget Sound *O. lurida*, with notable differences in the Oyster Bay population. In a reciprocal transplant study, Heare et al. (2017) found that Oyster Bay had considerably higher incidents of brooding, and reached maximum percent brooding 20-30 days earlier than Dabob Bay and Fidalgo Bay populations (145-159 degree days). Silliman et al. (2018) and Spencer et al. (2020) also found the Oyster Bay population to be the most reproductively active, and in Spencer et al. (2020) Oyster Bay oysters began releasing larvae on average 9.9 days earlier than Fidalgo and Dabob Bays (99 degree days earlier). Here, we find that Oyster Bay was the only population for which pCO_2_ exposure impacted reproductive processes, resulting in fewer females. A populations’ reproductive capacity is limited by the number of oysters spawning as females, therefore the productivity of oysters from Oyster Bay may be uniquely impacted by acidification.

If we look beyond our focal populations, individuals that grow slowly and are not highly fecund may be more capable of withstanding high pCO_2_ environments. Waldbusser *et al*. compared the response of *O. lurida* from Oregon with the faster growing Pacific oyster (*Crassostrea gigas*) to acidification, and suggested that slow shell secretion (a measure of growth rate) in *O. lurida* is a beneficial trait, contributing to their resilience to acidified conditions (Waldbusser et al., 2016). The growth rate of acidification-tolerant abalone is considerably lower than those sensitive to acidification, resulting in juveniles that are up to 80% smaller at 3-months old (Swezey et al., 2020). Whether growth rate is a proxy for resilience to acidification may not be applicable to all calcifiers, as faster growth is linked to ocean acidification resilience in selectively bred Sydney rock oysters (*Saccostrea glomerata*) (Parker et al., 2011; Stapp et al., 2018; Thompson et al., 2015). While no previous studies have associated reproductive investment with tolerance to acidification, a selectively bred line of Pacific oysters that are less sensitive to summer mortality and hypoxia also allocate less energy to reproductive tissue (Samain et al. 2007). Ultimately, the substantial and consistent resilience of the slow growing, moderately fecund Dabob Bay population in this study indicates that it is important to maintain a diversity of phenotypes, including those of lower fecundity and of smaller size. Oyster hatcheries routinely cull slow-growing larvae to maximize survival through metamorphosis, a practice that could be consistently removing stress-resilient genotypes. Highly fecund females and populations can also become overrepresented in cohorts of oyster seed. For long-term resilience to acidification and other stressors, commercial and restoration hatcheries should consider retaining slow-growing larvae and breed adults from a variety of sizes and productivity levels.

### Population-specific upregulation of some, but not all, detoxification genes in response to acidification

The transcriptional response of *O. lurida* acclimated to ocean acidification varied considerably by population, ranging from a robust response in Dabob Bay to no significant measurable response in Oyster Bay. The pronounced response of Dabob Bay to acidification could reflect that populations’ higher tolerance to stressors, and its ability to maintain altered homeostasis during the prolonged (52-day) exposure to acidification. For instance, Dabob Bay adults exposed to acidification contained a pronounced increase in transcripts from genes involved in detoxification, including several Cytochrome P450 (CYP), cytosolic sulfotransferases (SULT), and Glutathione transferases (GST). CYPs and SULTs are detoxification enzymes which metabolize both endogenous products (e.g. fatty acids, hormones) and xenobiotics (Coughtrie 2016; Snyder, 2000). Glutathiones are important cellular antioxidants which scavenge reactive metabolites, including hydrogen peroxide, and have been found at higher levels in many bivalves in response to acidification (Matozzo et al., 2013; Timmins-Schiffman et al., 2013), which may be related to their ability to protect proteins from oxidative stress (Abele et al., 2011; Sandamalika et al., 2019; Tomanek, 2015). Enhanced expression of GST and other intracellular stress mechanisms have similarly been observed in acidification-tolerant Sydney rock oysters upon intergenerational exposure to acidification (Goncalves et al., 2017), and direct exposure to thermal stress (McAfee et al., 2018). Two genes that codes for S-crystallins were also more prevalent in Dabob Bay oysters acclimated to high pCO_2_. S-crystallins are known to be structural components of cephalopod eyes, however they are apparently derived from GSTs (Tomarev & Piatigorsky, 1996) as they have very similar amino acid sequences, and while they are generally thought to lack enzymatic activity they are induced by heat shock (Lang et al., 2009). This could indicate that the Dabob Bay population activates additional detoxicant pathways for a more robust response to environmental stressors.

Interestingly, none of the typical enzymatic antioxidants (catalase, superoxide dismutase, peroxiredoxin) were differentially expressed in response to acidification in any population, despite detection and sufficient coverage. Opposing expression patterns have previously been reported for GST and enzymatic antioxidants in a transgenerationally acclimated acidification-tolerant line of Sydney rock oysters (Goncalves et al., 2016), which the authors suggested reflects distinct detoxification systems and/or stimuli. The glutathione scavenging system is also uniquely upregulated in a heat-tolerant species of vent polychaete compared to a heat-sensitive congener (Dilly et al., 2012). GST and related proteins may therefore act as the main cellular antioxidants in populations of oysters and other invertebrates tolerant to acidified conditions and other abiotic stressors.

### Population-specific changes in metabolism and energy production

Acidification altered expression of genes involved in lipid metabolism and transport in a population-specific pattern similar to the detoxification genes such that they ranged from highly upregulated in Dabob Bay to nearly unresponsive in Oyster Bay. These included genes that code for proteins in the peroxisome, which is an organelle that produces phospholipids, a major component of cell membranes, and metabolize long-chain fatty acids for energy production. The peroxisome proliferator-activated receptor gene, for instance, was much more active in the Dabob Bay population, which indicates the need for synthesis of peroxisome organelles. The co-activation of lipid metabolic and oxidation-reduction processes, most notably in Dabob Bay, could reflect enhanced mobilization of energy stores to support increased detoxicant synthesis (Goncalves et al., 2016; Hochachka & Somero, 2002; Mayor et al., 2015; Sokolova et al., 2012). There are previous reports of acidification-induced increases in intracellular energy production, particularly in organisms tolerant to acidification and following long-term exposure (reviewed by Strader et al., 2020), and coinciding with activation of antioxidant defenses (Goncalves et al., 2016).

Several other genes involved in mitochondrial energy production were altered in acidification-acclimated Dabob Bay oysters, suggesting increased energy production. This included the Acadl gene which codes for the Long-chain specific acyl-CoA dehydrogenase, a mitochondrial protein involved in production of energy from fats (specifically, fatty-acid beta-oxidation), and the si:dkey-18l1.1 gene (von Willebrand factor) which is involved in mitochondrial ATPase activity, and the alxA gene, which codes for alternative oxidase and which may increase mitochondrial respiration when the cytochrome respiratory pathway is restricted. There were also population-specific changes to a clustered mitochondria protein homolog (AAEL000794) which is involved in the distribution of mitochondria in the cytoplasm. Basal levels were fewer in Dabob Bay than other populations (not significantly), but were then substantially higher in Dabob Bay in high pCO_2_. The same expression pattern was observed in atad3-a (ATPase family AAA domain-containing protein 3-A), a protein that is essential for mitochondrial organization. Taken together, the basal and induced expression patterns in Dabob Bay could reflect a unique shift in mitochondrial efficiency and density in response to high pCO_2_ to produce sufficient energy to maintain cellular homeostasis in prolonged acidification exposure.

### Constitutive expression differences in the Dabob Bay population

We explored constitutive expression in Dabob Bay in an effort to understand why that population is unique in its transcriptional response to acidification, and how that might relate to its track record of high survival and stress tolerance (Heare et al., 2017; Silliman et al., 2018; Spencer et al., 2020). Many of the annotated genes that were uniquely expressed in control Dabob Bay adults were involved in immune functions (e.g. the complement system) and were enriched for antimicrobial and antiviral processes (e.g. cellular response to interferon-gamma), indicating that the Dabob Bay population is uniquely equipped to combat pathogens. Metabolic, growth, and reproduction processes were also overrepresented in Dabob Bay adult expression, and many of the same gene families were also uniquely expressed in Dabob Bay larvae. These annotated gene sets unique to Dabob Bay adults and larvae (220 and 25, respectively, Supplemental Materials) provide insight into the mechanisms behind the population’s unique energy distribution across life stages, and should be validated in other stress-tolerant, high-performing marine invertebrate populations.

Given its slow growth rate and relatively low/moderate fecundity, the Dabob Bay population was theorized to allocate more resources towards stress-response processes than other populations by constitutively expressing higher levels of genes that respond to acidified conditions. In this way, Dabob Bay would be uniquely “primed” for acidification by maintaining more transcripts of beneficial genes under typical conditions, which then become even more prevalent in acidification-acclimated oysters. Contrary to this prediction, those genes that were at higher levels in Dabob Bay constitutively become less abundant in response to acidification, and conversely those that were less abundant constitutively increased in acidification. The annotated genes that were depressed constitutively but were activated in response to acidification were largely involved in lipid transport and energy production, in addition to protein stabilization and cell migration. The annotated genes that were more active constitutively but then decreased in response to acidification were involved in immune function, cell cycle, and reproduction. Our findings suggest that populations of marine invertebrates tolerant to long-term acidification exposure are capable not necessarily because they are more prepared at the transcript level, but rather because they can mount then sustain a shift in resources to maintain lipid metabolic function, while down-regulating functions that are not critical to the acidification response, such as reproduction, immune function, and cell cycle.

### Population-specific carryover effect of parental exposure to acidification

Parental exposure to acidification resulted in larger larval offspring in the Oyster Bay population only. Oyster Bay larvae from parents exposed to acidification were on average 5µm (3%) larger than those from control parents. Larval size is positively associated with lipid content, growth rate, and feeding ability in many bivalves, and can reduce predation risk (Bailey, 1984; Gonzalez Araya et al., 2012; Helm et al., 1973; Marshall & Keough, 2007; Wilson et al., 1996). Increased larval size following intergenerational exposure to acidification and warming could therefore benefit some *O. lurida* populations in the wild, particularly those that also encounter environmental stressors (Gibbs et al., 2021). Our results align with previous studies showing that parental exposure to acidification can influence the physiology of invertebrate larval offspring. This was first detected in the Sydney rock oyster (Parker et al., 2012), as parental exposure to acidification resulted in higher larval survival and growth rates. Subsequently, some studies also reported positive intergenerational and transgenerational carryover effects (reviewed in Zhao et al., 2020), while others found negative effects (Parker et al., 2017; Venkataraman et al., 2019), or no signal of parental exposure (Clements et al., 2021). In the present study, carryover effects were only detected in one of three populations, indicating that parental priming may only be triggered by acidification in some genotypes or epigenotypes. Furthermore, populations unable to acclimatize directly to acidification (i.e. Oyster Bay in this study) may instead invest in parental priming.

The variety of intergenerational responses observed here and in previous studies could also stem from the mechanisms by which offspring are affected, which are theorized as changes to maternal provisioning, gamete mRNAs, and epigenetic changes (Eirin-Lopez & Putnam, 2019). If so, one would expect associated signals in larval transcriptomes, reflecting either energetic differences, remnant maternal mRNA transcripts, or regulatory shifts due to epigenetic changes (Gavery & Roberts, 2010, 2013). Curiously, despite size differences associated with parental treatments in the Oyster Bay population, there were no differences in gene expression. Differing sizes may therefore not be due to variable growth rate, as that would likely have been reflected in expression profiles (Meyer & Manahan, 2010; Pace et al., 2006). It is possible that cryptic intergenerational transcriptome plasticity could be induced by high pCO_2_ or other environmental stressors, but that was not revealed under ambient conditions in which larvae were reared. It remains unclear why intergenerational acidification exposure increases larval size in some populations and species.

### Puget Sound O. lurida populations have unique physiotypes which may be adaptive

To date, this and five other studies have characterized Puget Sound *O. lurida* with Fidalgo Bay, Dabob Bay, and Oyster Bay heritage (Heare et al., 2017; Heare et al., 2018; Silliman et al., 2018; Spencer et al., 2020; White et al. 2017). The three populations are close proximally (all are within the greater Puget Sound estuary), yet they represent distinct physiotypes in how they prioritize energy allocation constitutively, and when responding to an environmental stressor. Remarkably, in contrast to the other populations, high pCO_2_ elicited no transcriptional response in Oyster Bay, which raises the question as to where energy typically allocated towards reproduction and growth was utilized in Oyster Bay oysters. A previous study on the same populations provides insight into Oyster Bay population’s response. Heare et al. (2018) measured expression of targeted genes involved in the immediate (1-hr) stress-response following acute heat and mechanical shock, and reported that Oyster Bay was the only population for which a transcriptional response was detected. The discrepancy between Heare et al. (2018) and the present study could reflect unique responses to short-term vs. long-term abiotic stress. Specifically, those populations that are not capable of homeostatic stress-response over long periods could display the most pronounced short-term, acute response (i.e. Oyster Bay). We therefore suggest that the Oyster Bay population’s aerobic scope shifted nearer to or into the pessimus range compared to the other populations (Sokolova et al. 2012), and was not capable of maintaining its regulatory response for the full 52-day exposure to acidification. Time-series expression analysis would improve our understanding of populations’ varying abilities to respond to and maintain cellular functions over the course of prolonged exposure to acidification.

As suggested by Heare et al. (2017) and Silliman et al. (2018), the bay of origins’ distinct environments may explain varying physiotypes observed in Puget Sound *O. lurida*. Dabob Bay is located within the Hood Canal, which is a notoriously challenging environment for marine organisms. As a highly stratified, silled fjord with long residence times, it experiences slow turn-over, periods of hypoxia, and elevated temperature (Babson, Kawase, & MacCready, 2006; Banas et al., 2015; Khangaonkar et al., 2018; Newton et al., 2007). Thus, the slow-growing, transcriptionally responsive physiotype observed in Dabob Bay oysters may have arisen due to selection for genotypes that allocate a high proportion of energy to cellular maintenance and chronic stress response. Fidalgo Bay is located in the Puget Sound’s North Basin, and is more heavily influenced by tidal exchange and coastal oceanographic conditions. Fidalgo Bay characteristics (faster growth, larger mature size, low/moderate reproduction and transcriptional response) could reflect an ancestral population that has not experienced extreme selection events, and therefore represents a less specialized physiotype. Oyster Bay is located in Southern Puget Sound, which is a system of shallower finger-like basins that are highly productive, mixed, and experience large seasonal temperature swings (Moore et al., 2008). South Puget Sound is well suited for both wild and farmed shellfish, and may have preferentially selected for individuals that are highly fecund but lack the ability to acclimatize to acidification. Given that population-of-origin was also a dominant factor influencing gene expression in larvae, the genetic or epigenetic contributions to diverse physiologies should not be underestimated for populations of *O. lurida* and related species, even in small geographic scales.

## Conclusion

This is the first study to assess the transcriptional response of an oyster from the genus *Ostrea* to ocean acidification. In doing so, we greatly expand our understanding of how different oyster species will respond to shifting ocean conditions. There is increasing evidence that *Ostrea* spp. may be more tolerant than other oysters to acidification (Cole et al., 2016; Gray et al., 2019; Spencer et al., 2020; Waldbusser et al., 2016). Exploring cellular strategies in stress-tested larvae, a highly vulnerable stage, is an important avenue for future research. Furthermore, a physiological response spectrum was observed in three *O. lurida* populations exposed to ocean acidification. Given previous observations of stress tolerance in oysters from Dabob Bay we suggest that the robust transcriptomic changes in acclimated oysters and slow growth rate represent the more acidification-tolerant physiotype. However, observations of positive carryover in Oyster Bay indicates that intergenerational plasticity could improve the outlook for future generations of less tolerant physiotypes.

## Acknowledgements

Our gratitude to the following people who assisted with this project’s many daily tasks: Grace Crandall, Kaitlyn Mitchell, Olivia Smith, Megan Hintz, Rhonda Elliott, Lindsay Alma, Sam White, Hollie Putnam, Brent Vadopalas, Bayer and Jackson Roberts, my husband Ian and my mom Anne. This project would not be possible without the support and encouragement of the Puget Sound Restoration Fund, which housed the experiment and whose staff contributed expert knowledge of Olympia oyster reproduction and husbandry: Ryan Crim, Stuart Ryan, Alice Helker, Brian Allen, Betsy Peabody, Josh Bouma, and Joth Davis. Thank you to my committee members Rick Goetz, Jacqueline Padilla-Gamiño, Jennifer Ruesink, and Steven Roberts. This work was supported by the National Science Foundation Graduate Research Internship Program, the University of Washington College of the Environment Hall Conservation Research Award, and the National Shellfisheries Association Melbourne Carriker Award.

---

**Supplemental Table 1:**
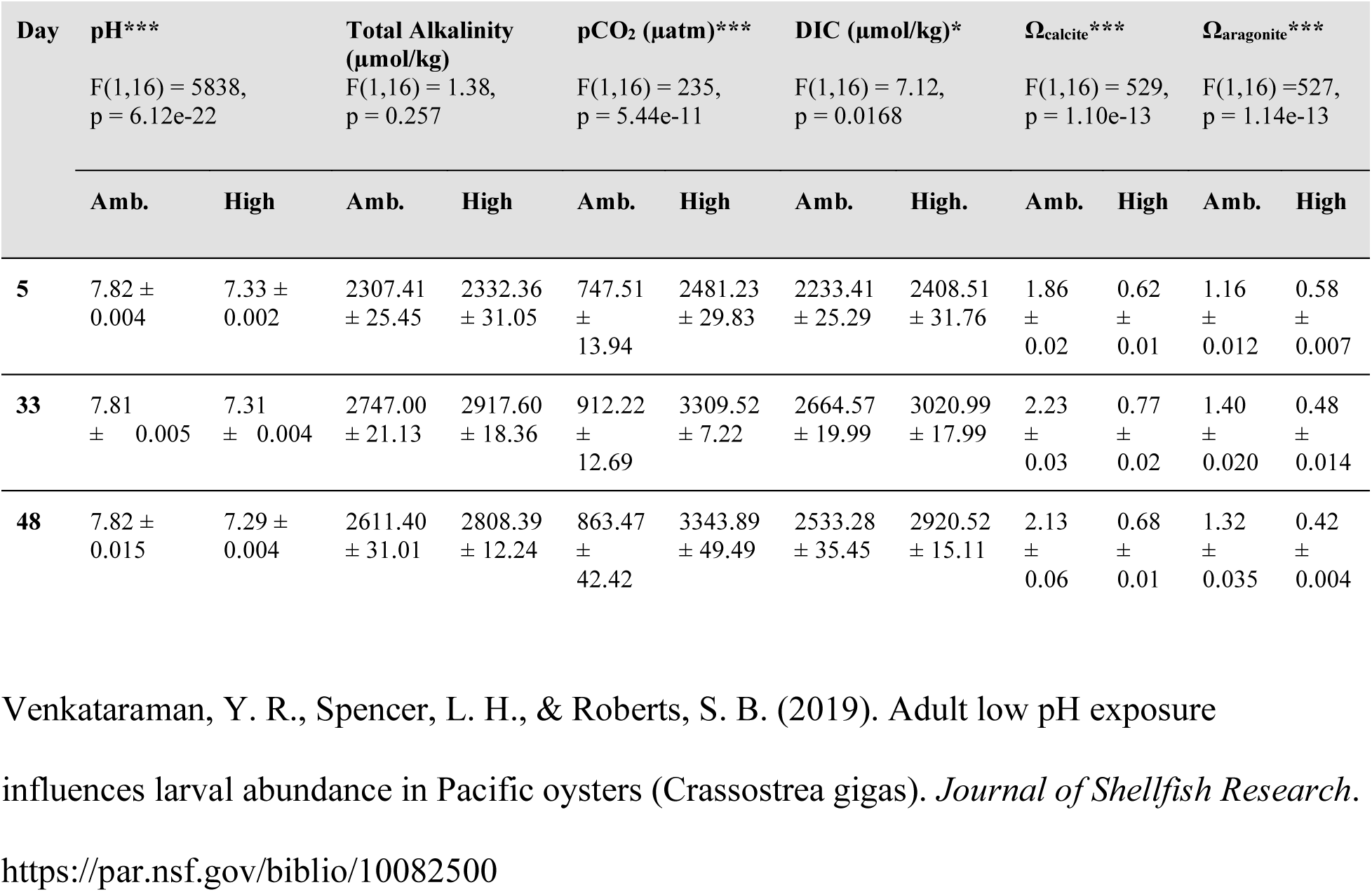
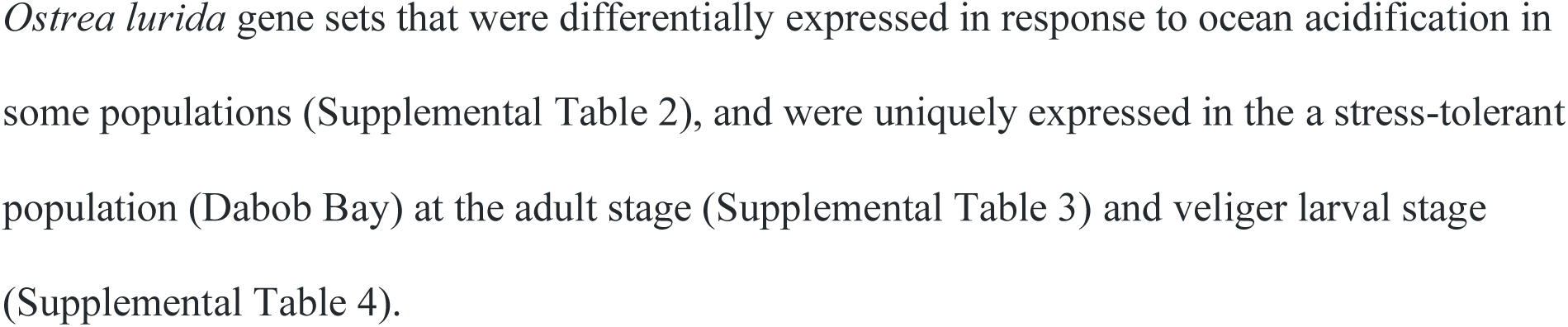
Carbonate chemistry parameters for three time points during the pCO_2_ treatments, which are averages (± SE) from three replicate tanks per treatment. All parameters except for total alkalinity differed significantly between control/ambient (Amb.) and experimental/high (High.) tanks (One-way ANOVA). More details are available in *Venkataraman et al., 2019*.

**Supplemental Table 2.**
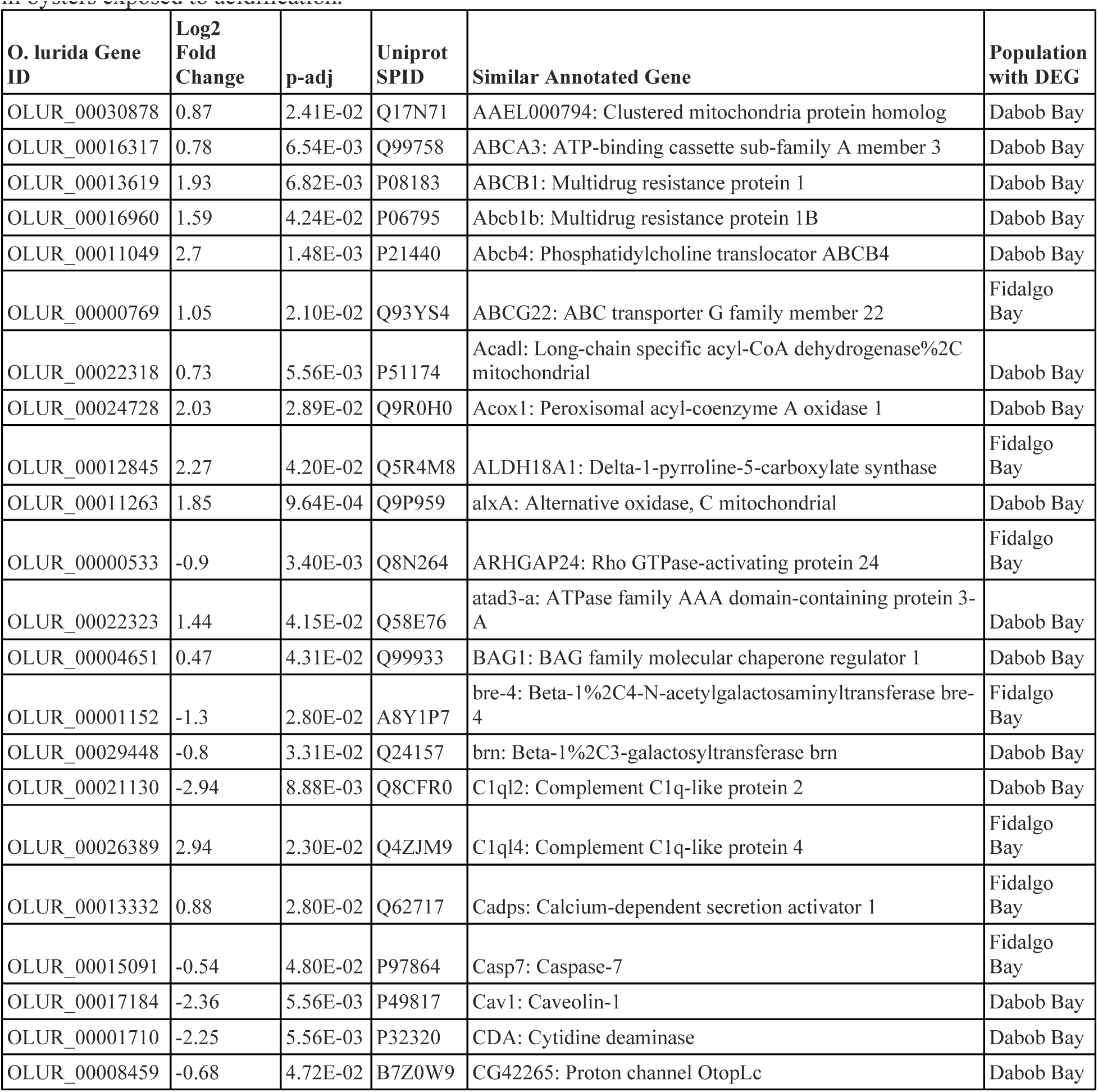
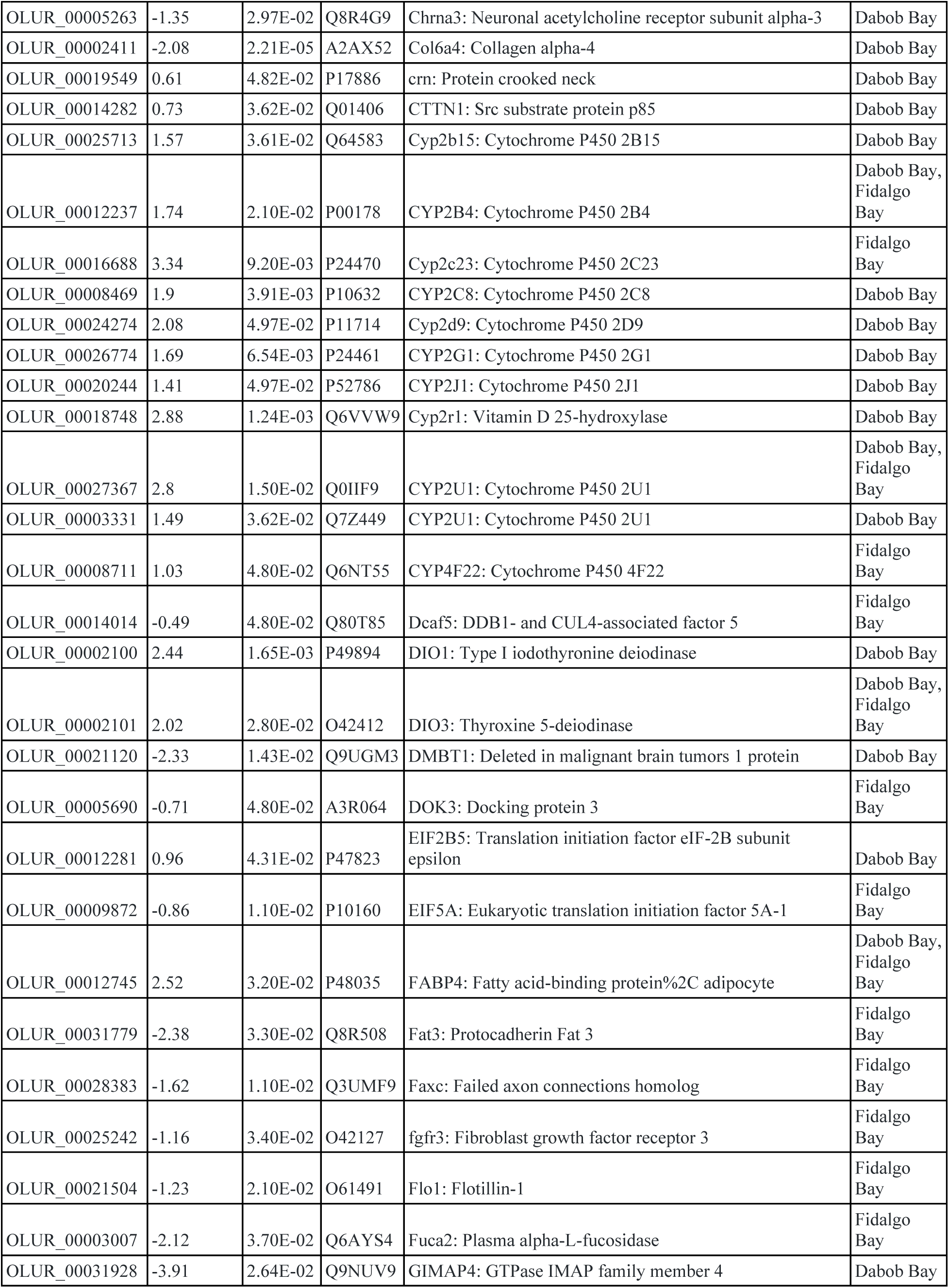
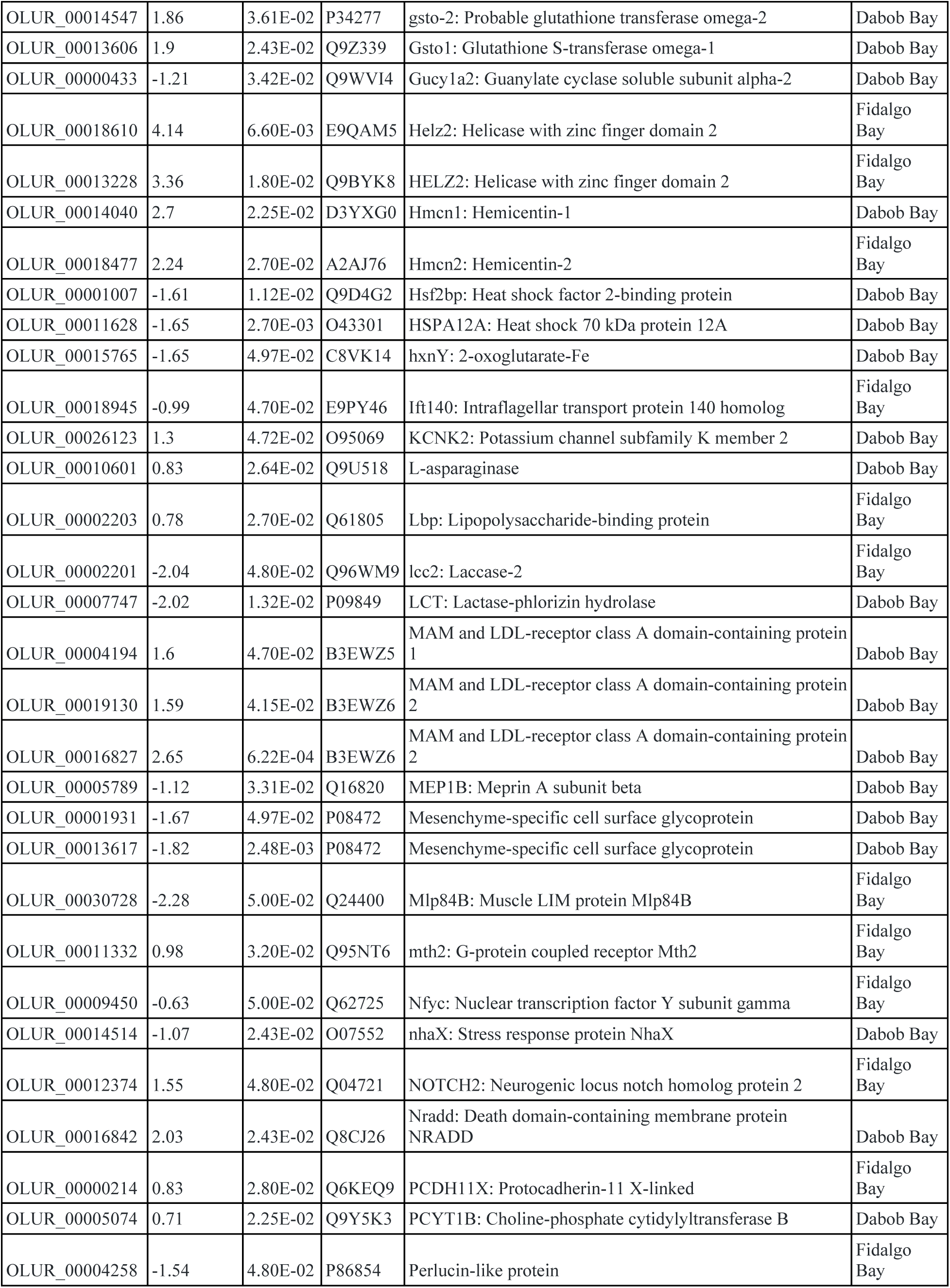
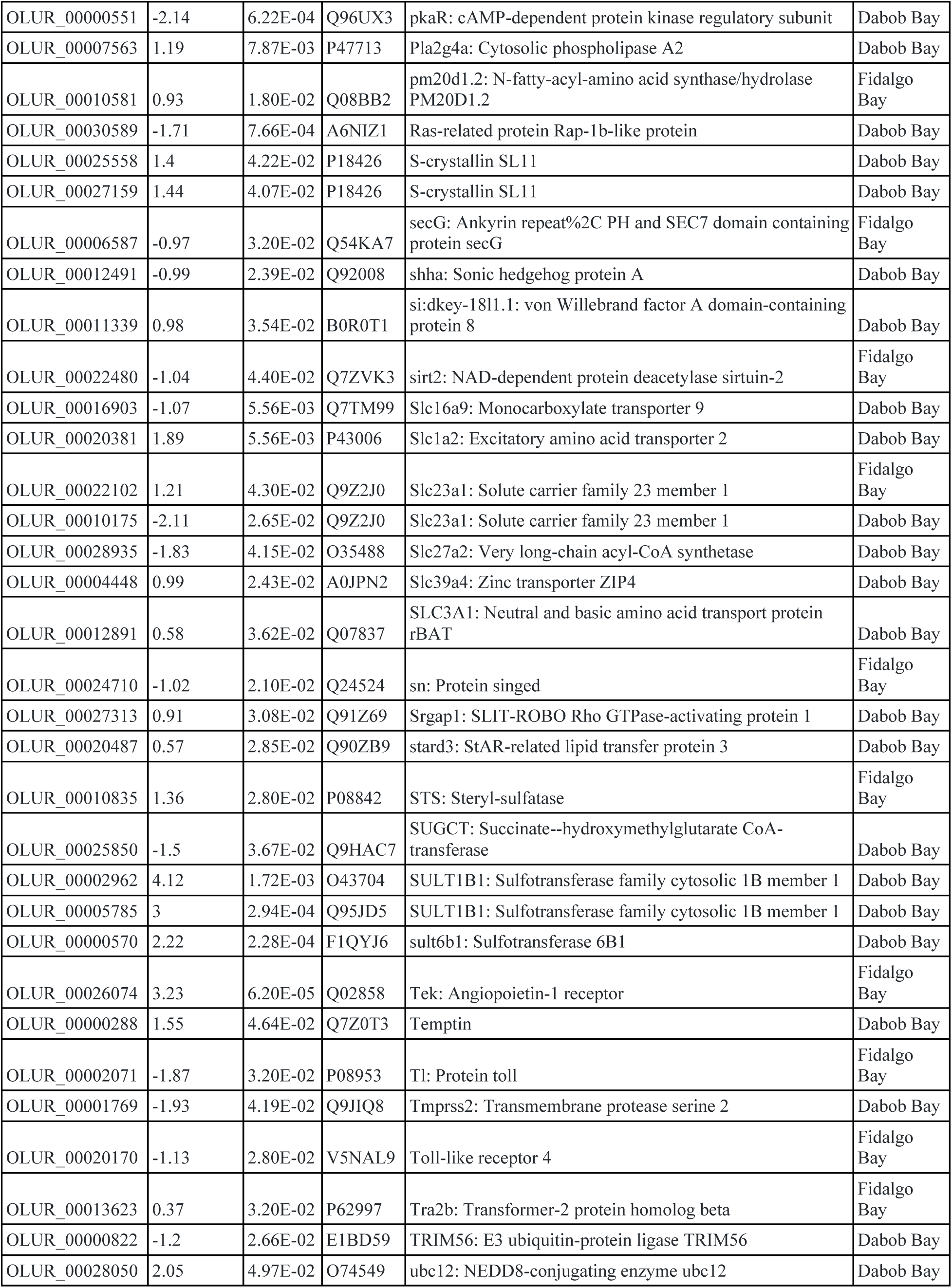
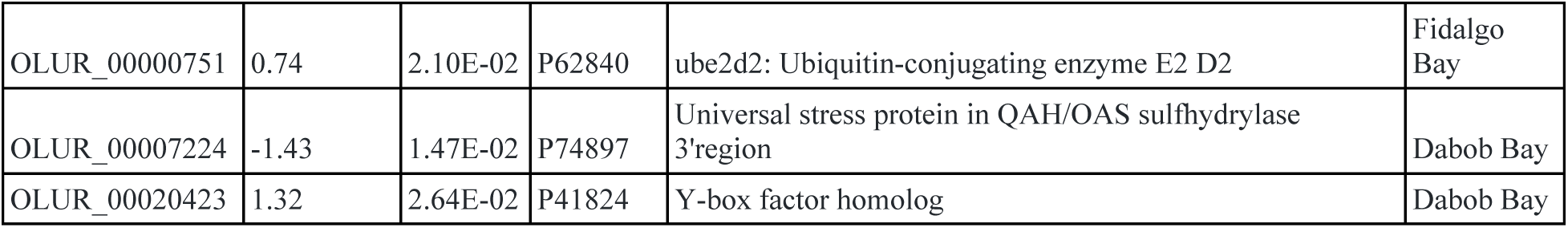
Genes that were differentially expressed in response to ocean acidification in *Ostrea lurida* populations, Dabob Bay and/or Fidalgo Bay. Positive Log2-FC values indicate abundances were higher in oysters exposed to acidification.

**Supplemental Table 3:**
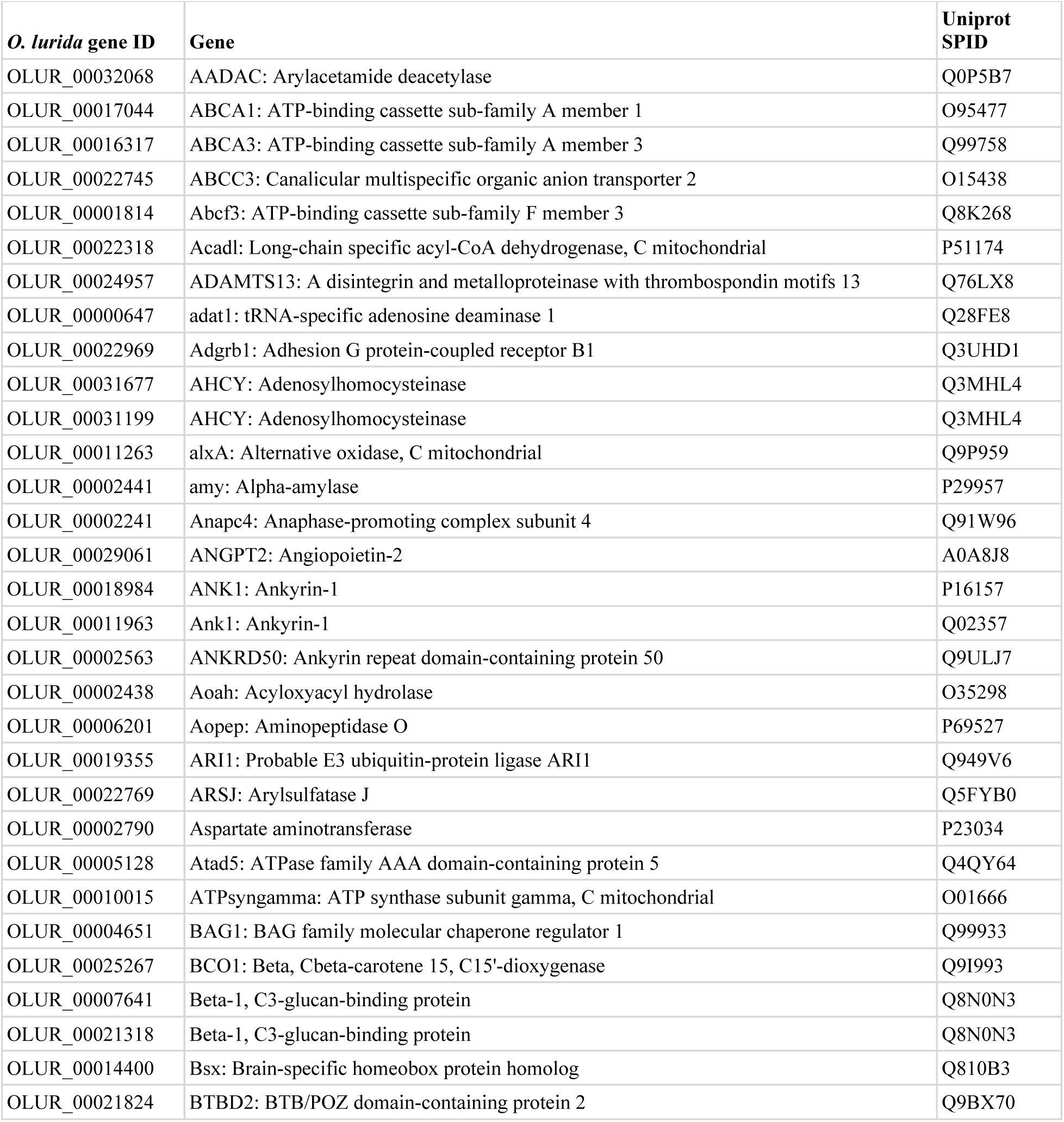
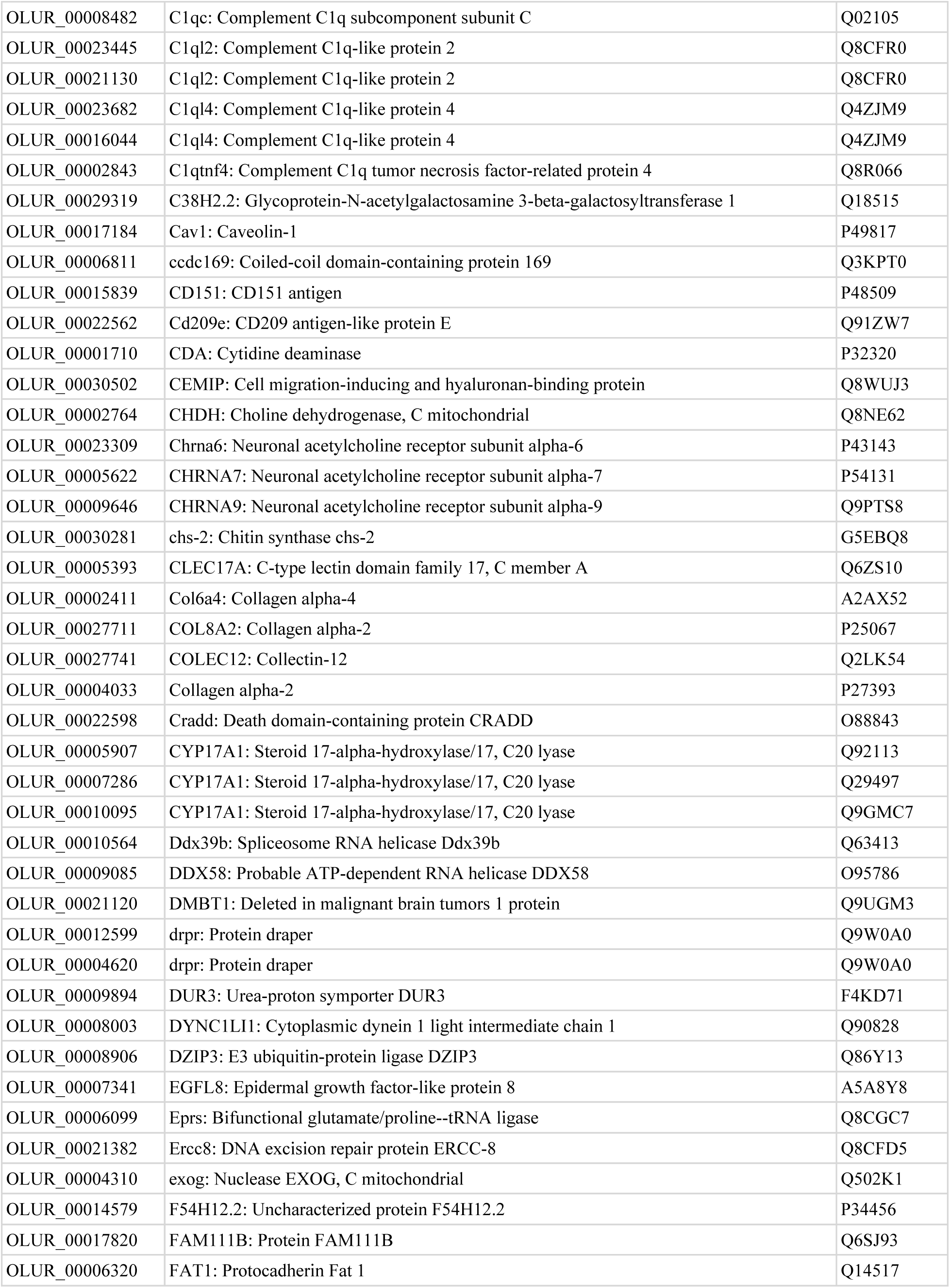
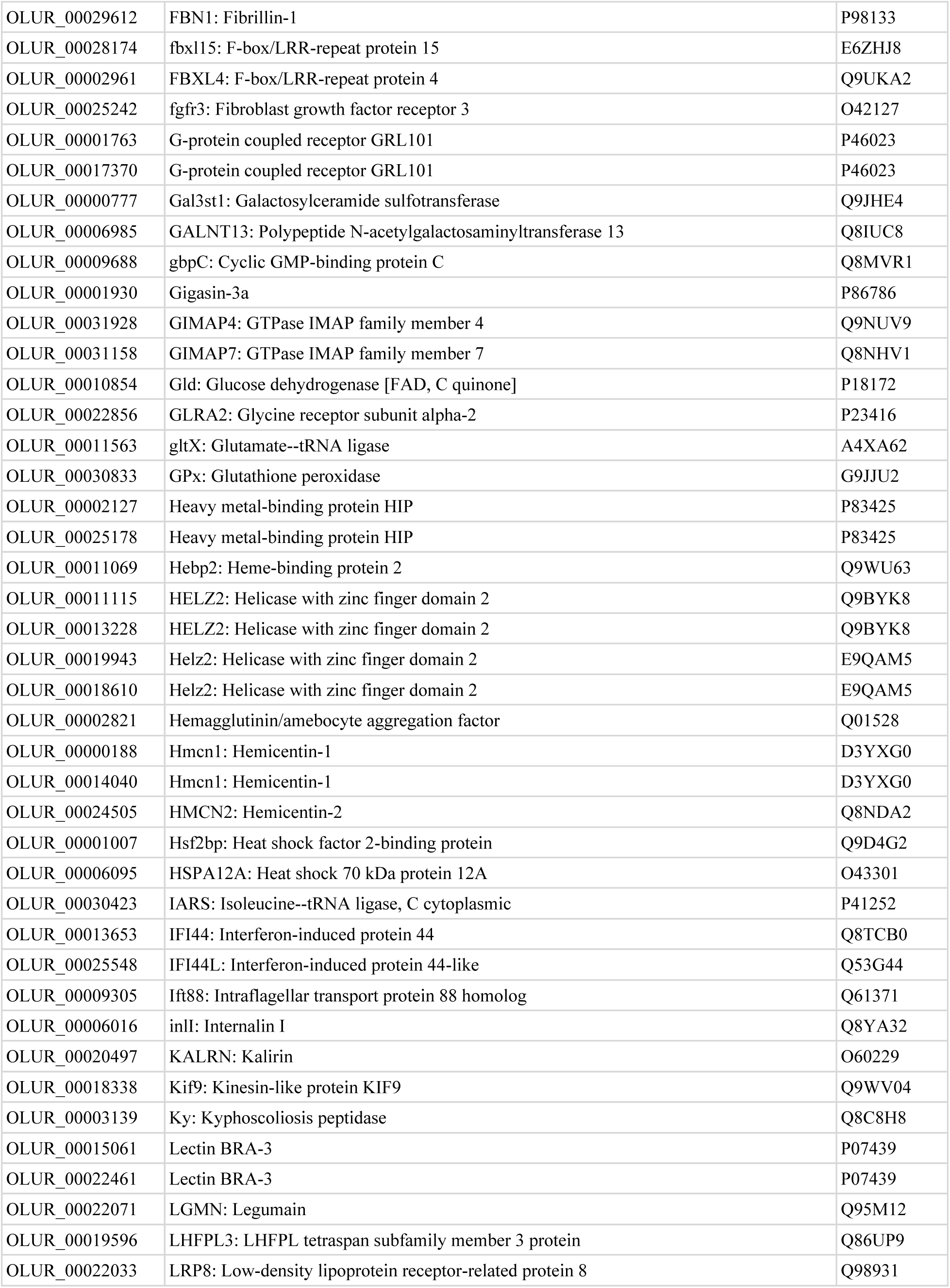
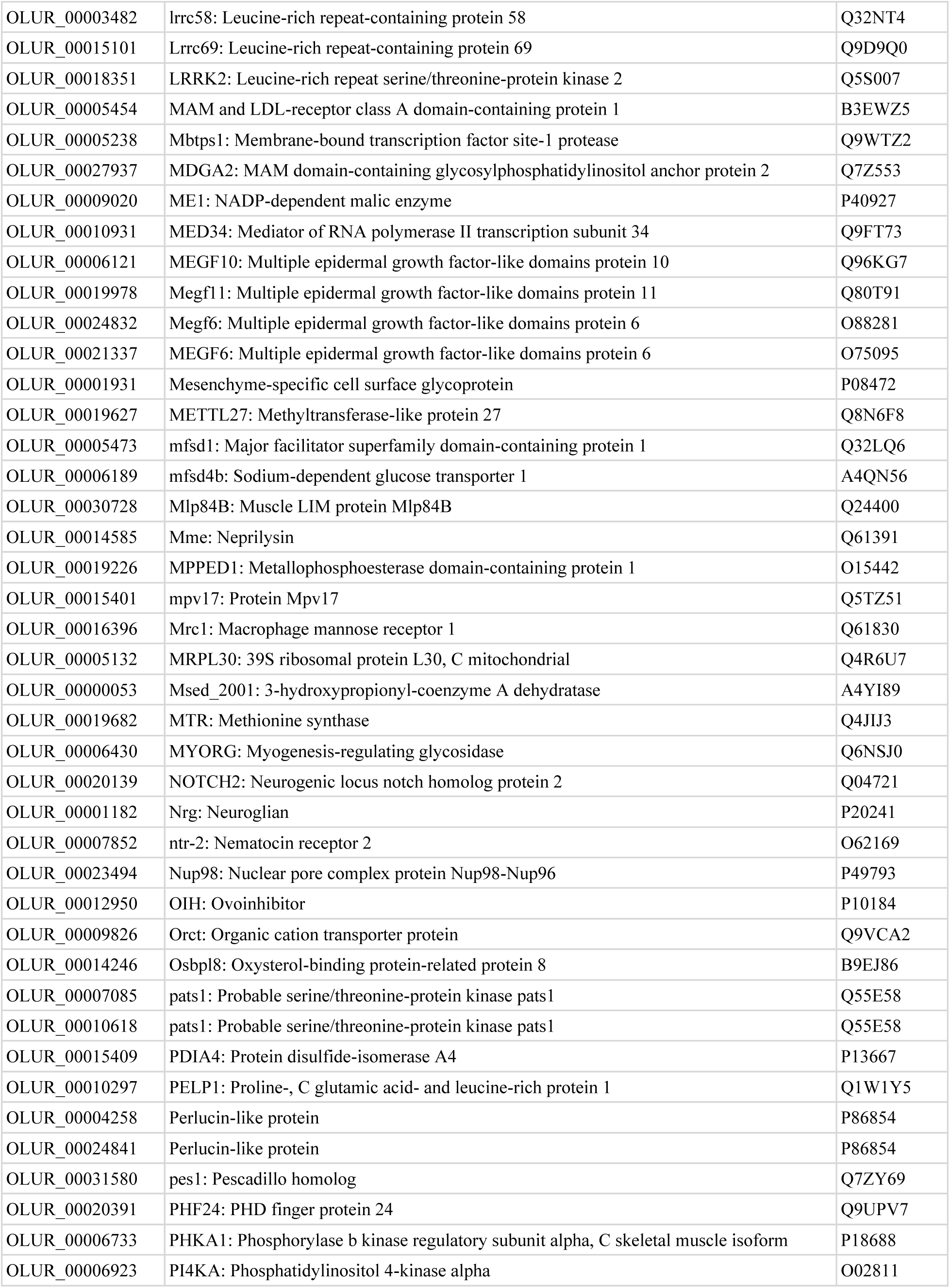
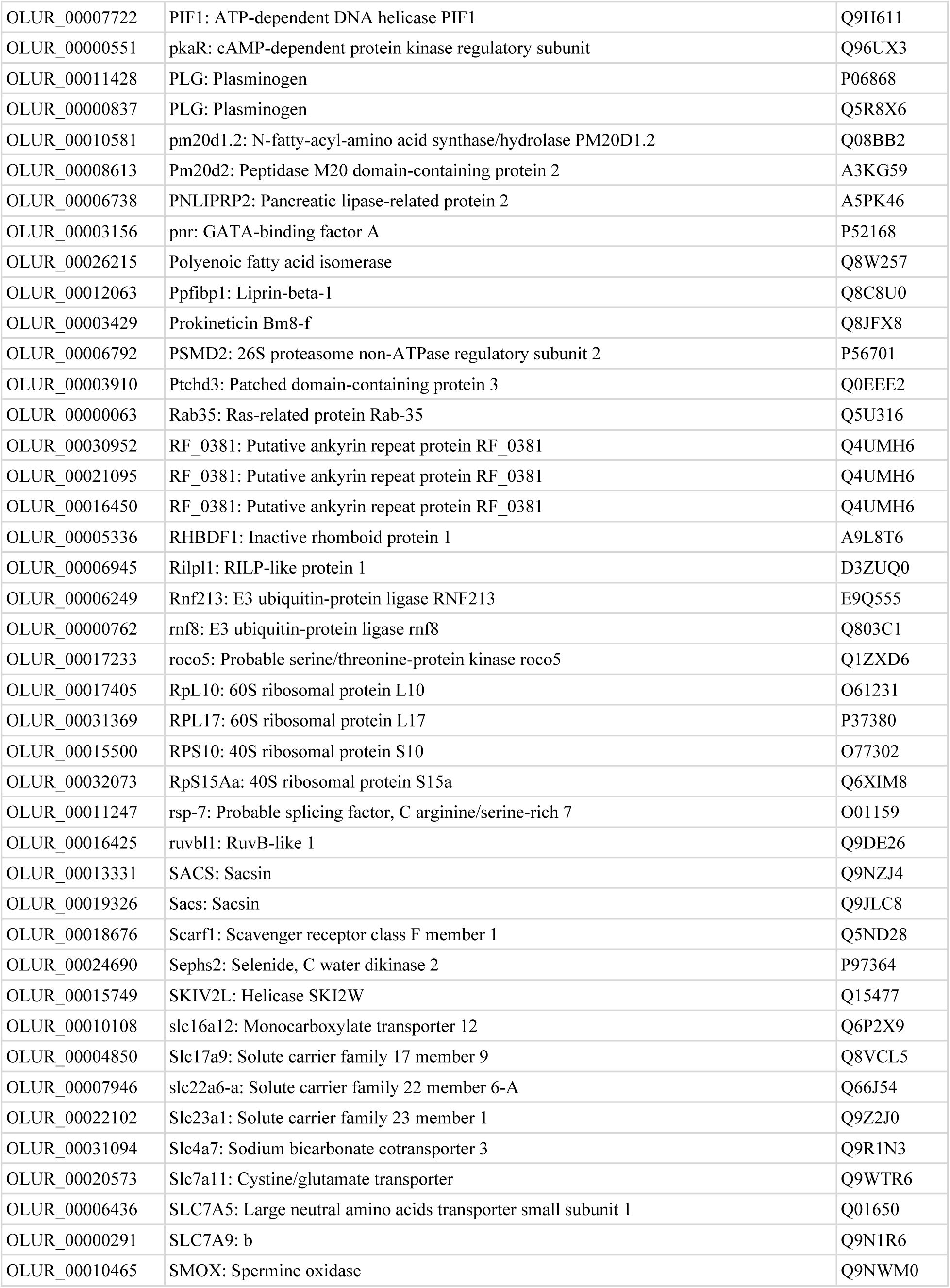
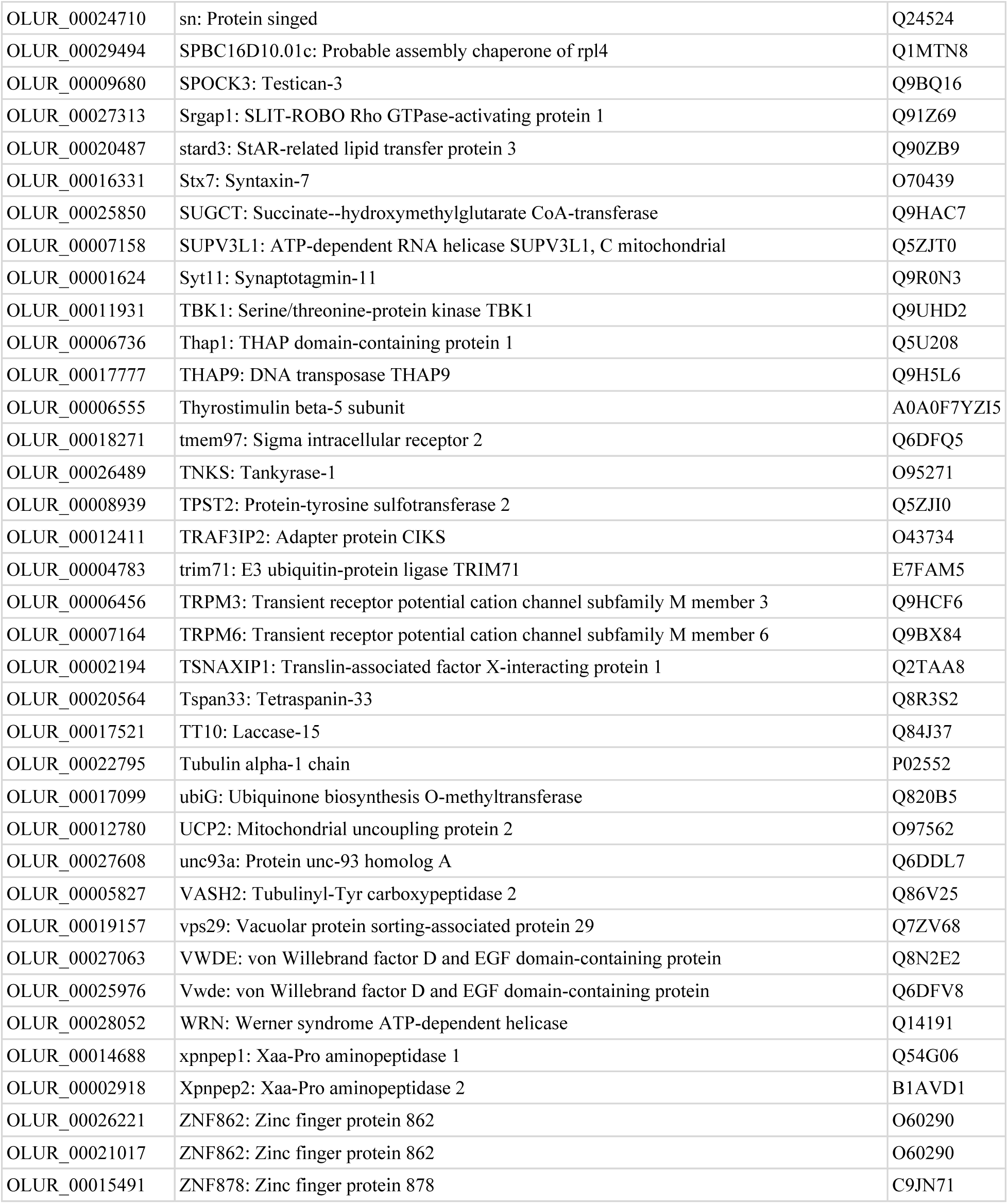
Genes that were constitutively expressed in Dabob Bay adults at different levels compared to other populations.

**Supplemental Table 4:** Genes that were constitutively expressed in Dabob Bay larvae at different levels compared to other populations.

| *O. lurida* gene ID | Gene | Uniprot SPID |
| --- | --- | --- |
| OLUR_00010887 | FMO5: Dimethylaniline monooxygenase [N-oxide-forming] 5 | P49326 |
| OLUR_00013658 | At5g10370: ATP-dependent RNA helicase DEAH1, 2C chloroplastic | F4KGU4 |
| OLUR_00013228 | HELZ2: Helicase with zinc finger domain 2 | Q9BYK8 |
| OLUR_00004343 | FAT2: Protocadherin Fat 2 | Q9NYQ8 |
| OLUR_00016966 | DDB_G0292642: Uncharacterized protein DDB_G0292642 | Q54CX4 |
| OLUR_00022057 | MSMEG_3950: Universal stress protein MSMEG_3950/MSMEI_3859 | A0QZA1 |
| OLUR_00025230 | ghrA: Glyoxylate/hydroxypyruvate reductase A | Q8FIT1 |
| OLUR_00000611 | L-proline trans-4-hydroxylase | R9UTQ8 |
| OLUR_00029178 | ucpB: Mitochondrial substrate carrier family protein ucpB | B0G143 |
| OLUR_00004362 | bmp2: Bone morphogenetic protein 2 | Q804S2 |
| OLUR_00019157 | vps29: Vacuolar protein sorting-associated protein 29 | Q7ZV68 |
| OLUR_00031354 | DDB_G0287015: TM2 domain-containing protein DDB_G0287015 | Q54KZ0 |
| OLUR_00003900 | AAO: L-ascorbate oxidase | Q40588 |
| OLUR_00004324 | UNC93A: Protein unc-93 homolog A | A2VE54 |
| OLUR_00003482 | lrrc58: Leucine-rich repeat-containing protein 58 | Q32NT4 |
| OLUR_00013735 | cep290: Centrosomal protein of 290 kDa | P85001 |
| OLUR_00018610 | Helz2: Helicase with zinc finger domain 2 | E9QAM5 |
| OLUR_00019426 | K02F3.12: Putative ATP-dependent DNA helicase Q1 | Q9TXJ8 |
| OLUR_00022209 | Ectin (Fragment) | B3EWZ8 |
| OLUR_00020882 | UBA6: Ubiquitin-like modifier-activating enzyme 6 | A0AVT1 |
| OLUR_00005664 | ABCC1: Multidrug resistance-associated protein 1 | Q5F364 |
| OLUR_00001519 | HET-E1: Vegetative incompatibility protein HET-E-1 | Q00808 |
| OLUR_00025496 | HSPA12B: Heat shock 70 kDa protein 12B | Q96MM6 |
| OLUR_00019143 | NADSYN1: Glutamine-dependent NAD(+) synthetase | Q5ZMA6 |
| OLUR_00010983 | CLEC3A: C-type lectin domain family 3 member A | O75596 |

## Notes

### Competing Interest Statement

The authors have declared no competing interest.

